# Individualized Gaussian Process-based Prediction of Memory Performance and Biomarker Status in Ageing and Alzheimer’s disease

**DOI:** 10.1101/2022.03.14.484226

**Authors:** A. Nemali, N. Vockert, D. Berron, A. Maas, R. Yakupov, O. Peters, D. Gref, N. Cosma, L. Preis, J. Priller, E. Spruth, S. Altenstein, A. Lohse, K. Fliessbach, O. Kimmich, I. Vogt, J. Wiltfang, N. Hansen, C. Bartels, B.H. Schott, F. Maier, D. Meiberth, W. Glanz, E. Incesoy, M. Butryn, K. Buerger, D. Janowitz, M. Ewers, R. Perneczhy, B. Rauchmann, L. Burow, S. Teipel, I. Kilimann, D. Göerß, M. Dyrba, C. Laske, M. Munk, C. Sanzenbacher, S. Müller, A. Spottke, N. Roy, M. Heneka, F. Brosseron, S. Roeske, L. Dobisch, A. Ramirez, M. Ewers, P. Dechent, K. Scheffler, L. Kleineidam, S. Wolfsgruber, M. Wagner, F. Jessen, E. Duzel, G. Ziegler

**Author notes:** Corresponding author at: Institute of Cognitive Neurology and Dementia Research, Otto-von-Guericke University Magdeburg, Germany., E-mail address (A. Nemali). The two authors contributed equally to this paper (E. Duzel, G. Ziegler).

## Abstract

Neuroimaging markers based on Magnetic Resonance Imaging (MRI) combined with various other measures (such as informative covariates, vascular risks, brain activity, neuropsychological test etc.,) might provide useful predictions of clinical outcomes during progression towards Alzheimer’s disease (AD). The Bayesian approach aims to provide a trade-off by employing relevant features combinations to build decision support systems in clinical settings where uncertainties are relevant. We tested the approach in the MRI data across 959 subjects, aged 59-89 years and 453 subjects with available neuropsychological test scores and CSF biomarker status (amyloid-beta (*Aβ*)42/40 & and phosphorylated tau (pTau)) from a large sample multi-centric observational cohort (DELCODE). In order to explore the beneficial combinations of information from different sources, we presented a MRI-based predictive modelling of memory performance and CSF biomarker status (positive or negative) in the healthy ageing group as well as subjects at risk of Alzheimer’s disease using a Gaussian process multikernel framework. Furthermore, we systematically evaluated predictive combinations of input feature sets and their model variations, i.e. (A) combinations of brain tissue classes and feature type (modulated vs. unmodulated), choices of filter size of smoothing (ranging from 0 to 15 mm full width at half maximum), and image resolution (1mm, 2mm, 4mm and 8mm); (B) incorporating demography and covariates (C) the impact of the size of the training data set (i.e., number of subjects); (D) the influence of reducing the dimensions of data and (E) choice of kernel types. Finally, the approach was tested to reveal individual cognitive scores at follow-up (up to 4 years) using the baseline features. The highest accuracy for memory performance prediction was obtained for a combination of neuroimaging markers, demographics, genetic information (ApoE4) and CSF-biomarkers explaining 57% of outcome variance in out of sample predictions. The best accuracy for *Aβ*42*/*40 status classification was achieved for combination demographics, ApoE4 and memory score while usage of structural MRI improved the classification of individual patient’s pTau status.

## 1. Introduction

Alzheimer’s disease (AD), a most common cause of dementia, is a progressive neurodegenerative disease affecting the ageing population worldwide. AD predominantly impairs the memory domain, eventually leading to the state of dementia where most cognitive functions are impaired and daily life activities are disrupted. (Gaugler et al., 2019). The current understanding of the disease pathology states that AD progression starts long before the clinically manifest dementia (Morris, 2005; Jack et al., 2016; Dubois et al., 2016; Beason-Held et al., 2013). According to the well-established Amyloid-Cascade-Hypothesis, AD begins with amyloid-beta (*Aβ*) protein accumulation and tau pathology followed by neurodegeneration (atrophy of the neuropil and loss of neurons causing brain atrophy) and cognitive impairment (Murphy and LeVine III, 2010; Jack Jr et al., 2013a; Blennow et al., 2012). Neurodegeneration may be characterized by measuring total tau in CSF, using magnetic resonance imaging (MRI) techniques, by assessing local brain volumes (Frisoni et al., 2010; Knešaurek, 2015; Besson et al., 2015), and metabolism using positron-emission tomography (PET) (Forsberg et al., 2008; Humpel, 2011).

An important challenge in the clinical practice as well as in AD research is to determine the relationship between cognitive impairment, structural brain alterations, and biomarkers of amyloid or tau pathology, both cross-sectionally and with regards to longitudinal cognitive decline. Here, we address some of these challenges from the perspective of predictive modelling and the machine learning approach.

Scientifically, this approach can reveal mechanisms contributing to cognitive or brain reserve (Stern et al., 2020). Given the regional distribution of amyloid and tau pathology in AD, it is likely that the volume of the affected brain regions are predictive of biomarker burden (Ossenkoppele et al., 2016). To the extent that the same brain regions are also components of neurocognitive circuits such as for memory formation, they predict both biomarker status and cognitive performance cross-sectionally. If, on the other hand, MR-based volumes and demographic variables jointly predict substantially more variance in cognitive performance than CSF biomarkers, this might indicate the joint operation of cognitive and brain reserve mechanisms. From a clinical perspective, understanding the relative contribution of demographics/genetic variables (ApoE4), CSF biomarker status and patterns of brain volumes to predict cognitive decline is relevant for early detection and diagnosis. For instance, if regional brain volumes substantially add to the prediction of longitudinal cognitive decline, over and above demographic/genetic variables and biomarker status, MRI should become part of the decision making process for identifying rapid decliners and thereby of individuals who should be prioritised for disease-modifying treatments (Cummings et al., 2016).

Machine learning (ML) or artificial intelligence based on computational models and large datasets has boosted medical image analysis and clinical decision support over the past decade (Davatzikos et al., 2019; Mateos-Pérez et al., 2018; Arbabshirani et al., 2017; Salvatore et al., 2016). It is a powerful and promising set of tools to assist doctors with quantitative evidence about individual patients, by supporting diagnosis and prognosis of developmental, psychiatric and neurodegenerative disorders. These pattern-based predictions can be particularly useful in subjects at risk of dementia, to obtain a clinical staging for potential treatments and also introduce more precise diagnostic and predictive biomarkers of clinical outcomes, to achieve an informed matching of available treatments to patients (Rathore et al., 2017).

There are conceptually two different types of ML approaches, one is to predict continuous variables (known as regression) and the other one is to predict binary variables *y* ∈ {−1, 1} (known as classification) (Bishop, 2006). One particularly important application of ML is to implement a predictive model for biomarkers or cognitive decline. Studies using regression for prediction of cognitive performance differences have been previously introduced for assessing cognitive competence/ability and predicting IQ in healthy individuals (Rohde and Thompson, 2007; Bradley and Caldwell, 1980; Barber, 2005). One particularly important application of ML is to implement a predictive model for biomarkers or cognitive decline. Similarly, studies predict cognitive performance (Kandiah et al., 2014; Doraiswamy et al., 1998; Woodard et al., 2010) and biomarkers status (Prestia et al., 2015; Besson et al., 2015) in older cognitively impaired individuals. Two aspects of effective predictions are to incorporate multiple sources of information such as different morphometric brain properties e.g. from different features or tissue classes (Monté-Rubio et al., 2018), image modalities (e.g. T1 and FLAIR) (Amyot et al., 2015; Johnson et al., 2012; Sui et al., 2012), and accounting for demographic background and subjects-specific covariates (Ziegler et al., 2014). In recent years, many new tools for predicting cognitive performance and biomarkers status of AD have been integrated and support individualized diagnosis and prognosis (Marquand et al., 2014; Davatzikos et al., 2001; Franke et al., 2010; Rathore et al., 2017). For classification and regression, in addition to classical kernel-based methods such as support vector machines (Shawe-Taylor and Cristianini, 2004), Gaussian processes (Rasmussen and Williams, 2006), multi-kernel approaches (Pettersson-Yeo et al., 2014; Aksman et al., 2019), and neural networks (Fisher et al., 2019; Jo et al., 2019; Dyrba et al., 2021) are increasingly applied.

Machine learning using the so-called ‘kernel-method’ has been shown to reveal powerful applications in the context of neuroimaging (Dosenbach et al., 2010). A kernel serves as a bottleneck in learning algorithms that captures (and compresses) relevant individual differences even for high-dimensional input data such as images (Shawe-Taylor et al., 2004). Furthermore, kernel methods have shown promising performance in studies comparing different machine learning frameworks (Jollans et al., 2019) and even similar performance, when compared to the recent deep neural networks (He et al., 2020). Here we generalize ideas from (Ashburner and Klöppel, 2011) for multivariate classification and regression based on (primarily) linear kernels efficiently representing different morphometric brain features (such as gray matter (GM), white matter (WM) & cerebrospinal fluid (CSF); see also (Monté-Rubio et al., 2018)), and subjects-specific covariates (for-example age, sex & education; see also (Ziegler et al., 2014)). Using kernels in combination enables to explore the contribution of information from different sources (for-instance morphometric brain features & subject-specific covariates) (Zu et al., 2016). Previous studies described how to combine multiple kernels (Rakotomamonjy et al., 2008; Gönen and Alpaydın, 2011; Bach et al., 2004) and report that usage of multiple kernels can improve overall predictive performance (Gönen and Alpaydın, 2011; Wilson and Adams, 2013).

There is a compelling argument towards a Bayesian treatment of uncertainties of predictions in clinical and translational applications (Kim and Ghahramani, 2006; Filippone et al., 2012). We therefore, focus on Gaussian Process (GP) models, which enable full probabilistic predictions and Bayesian model comparisons (Rasmussen and Williams, 2006). GPs are flexible tools for regression of continuous metrics (such as current and memory performance at follow-up) and classification of binary variables (such as biomarker positivity). Moreover, GPs allow effective handling of multiple information sources of individual differences using multi-kernel learning (Marquand et al., 2014; Rakotomamonjy et al., 2008; Bach et al., 2004; Zu et al., 2016).

In this study, we present an application of multi-kernel learning to predict the cognitive performance and the biomarker status of subjects in a large well-characterized longitudinal MRI cohort of ageing and AD subjects. The applied GP model for prediction of cognitive performance and biomarker status determines the optimal (positive) weighted combination of image-based kernel matrices. Using the optimally selected kernels, we first assess the influence of (A) using alternative combinations of kernels from different morphometric aspects such as tissue classes; (B) using different spatial feature resolutions & feature types (modulated vs un-modulated); (C) the importance of incorporating demography and covariates; (D) impact of training size; (E) influence of reducing the dimensions of MR data and (E) non-linear kernels; all these aspects are evaluated concerning predictive performance (marginal likelihood). Finally, the clinical utility of imaging metrics will be assessed by their predictive power for new data samples including longitudinal cognitive follow-up data using 10-fold nested cross-validation.

## 2. Methods

### 2.1. Predictive model

A predictive model of cognitive ageing accurately predicts target (or output) variable *y*^*^ such as individual memory performance of an older study participant based on a new input test data sample **x**^*^ such as an MRI scan and/or subject-specific covariates. This model can be implemented by learning some unknown function *f* that maps input data features to outputs using a large set of training data 𝒟= {**X, y}**, where **X** represents input data of the training sample of MR features and a set of informative covariates e.g. [*age, sex, education*] obtained from concatenating individual features **x**_*i*_ for all *n* training subjects. In this work, we focus on two clinically relevant prediction scenarios. First, we aim at an MR-based prediction of individual memory performance (at baseline or follow-up) where all targets *y*_*i*_ (for subject *i*) refer to observations of a continuous variable. Secondly, using similar MRI feature sets we aim to classify subjects regarding their biomarker status i.e. *y*_*i*_ ∈ {−1, +1}. In the following section, we briefly revisit regression of continuous variables, and binary classification using multi-kernel learning and Gaussian processes.

### 2.2. Gaussian process regression

In general a Gaussian process (GP) describes a (prior) distribution over functions, which is fully specified by its mean *m* and covariance *k* (for a technical introduction see Rasmussen and Williams, 2006)

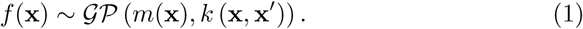

Gaussian process regression (GPR) is a non-parametric generalization of linear regression and can be described for training data 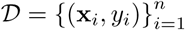 as

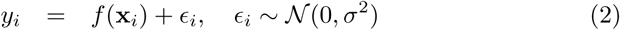

with subject index *i*, and an additive independent identically distributed Gaussian noise with variance *σ*^2^. In this study the latent function *f* (**x**) incorporates our knowledge about an older participant’s memory score in different locations **x** of the brain morphometric and covariate feature space describing individual differences. More specifically the prior mean and covariance are functions of the input data and imply a prior distribution over latent mappings *f*

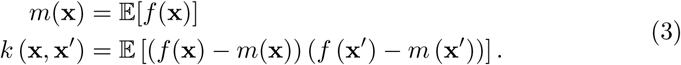

The principal idea of GPR then is that we assume that the covariance of cognitive scores *f*(**x**) is a function of the similarities of participants’ brain morphometry, expressed by a specific choice of a kernel mapping *k*. In this particular study, we further explored the following choices for *k* using linear kernels, squared exponential (SE) kernels for imaging modalities, and automatic relevance determination (ARD) for small dimensional covariate spaces:

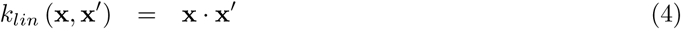

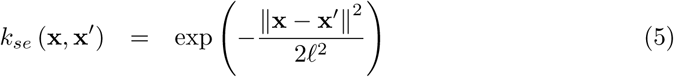

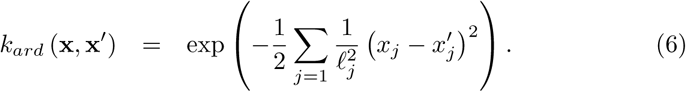

When using SE kernels and ARD we introduce additional hyperparameters such as characteristic length scale 𝓁_*j*_ of feature dimension *j*. Having usually noisy observations the above model implies the following covariance for observed memory scores:

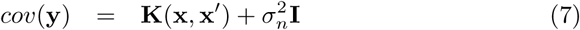

with **y** referring to a column vector of all observed memory scores and **K**(**x, x**′) denoting the evaluated kernel for all pairs of training points **x**. Furthermore we are interested in predictive distributions for new test sample **x**^*^ given a large dataset. This can be obtained using joint distribution of training **y** and testing **y**^*^ samples, and then conditioning **y**^*^|**y** gives us predictive mean and variance

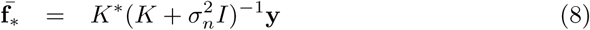

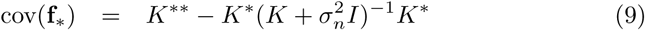

using *K* = **K**(**x, x**), *K*^*^ = **K**(**x, x**^*^) and *K*^**^ = **K**(**x**^*^, **x**^*^) for notation simplicity. The robustness of the GPR model is dependent on the choice of covariance function and its parameters *θ*. In order to choose sensible parameter estimates, we introduce the expression for the model evidence (or marginal likelihood) given by

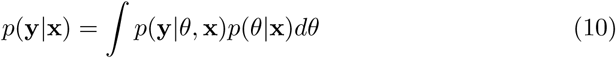

using the likelihood *p*(**y** |*θ*, **x**) and the prior *p*(*θ*|**x**) ∼ 𝒩(**0**, *K*). Integrating over factorized Gaussian likelihood and prior and evaluating the terms further results in the log marginal likelihood

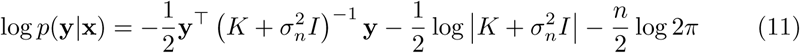

which is further used for model training and optimization of the (hyper-) parameters of the GP model. For efficient computation and numerical stability of matrix inversion, we use Cholesky decomposition (Rasmussen and Williams (2006)).

### 2.3. Gaussian process classifier

A Gaussian process classifier (GPC) for binary classification, *y*_*i*_ ∈ {−1, 1} is a non-parametric generalization of linear logistic regression. It places a GP prior over the latent function *f* (*x*) and maps it through the logistic function to model posterior class probability as *p*(*y* = 1| **x**) = *σ*(*f* (**x**)) (for details see Rasmussen and Williams, 2006). The inference of GPC is a two-step process. The first step is similar to GPR, which involves to compute the latent variable predictions 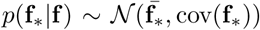 for the test sample similar to eq. (8) and eq. (9). In the second step, the latent function **f**_*_ is squeezed through a sigmoid function to estimate class membership probability given by

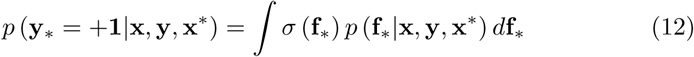

accounting for all uncertainties. However in contrast to GPR eq. (12) is not tractable analytically for certain sigmoid functions (such as the logistic sigmoid) and therefore we make use of the Laplace approximation to compute the approximation of the latent variable distribution

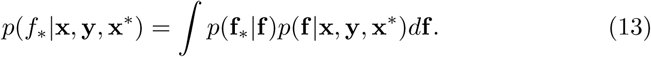

GPC model optimization of covariance functions and its parameters *θ* can be performed similarly to GPR but using an approximation of the marginal likelihood. For more details about Laplace approximation can be found elsewhere (Bishop, 2006; Rasmussen and Williams, 2006).

### 2.4. Multi-kernel learning

In order to enable the contribution of multiple imaging modalities (or feature types) and covariates to the predictions of individual memory performance and bio-marker status, we apply so called multi-kernel learning using a linear combination of kernels (Pettersson-Yeo et al., 2014; Aksman et al., 2019; Marquand et al., 2014)

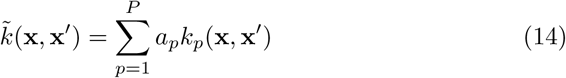

with a-priori unknown amplitude (hyper-) parameters [*a*_1_, …, *a*_*p*_] for p feature types, such as GM volume, WM volumes extracted from T1 weighted MRI scans and covariates such as age. *k*_*p*_ can be chosen using above choices for kernels (*k*_*lin*_, *k*_*se*_ or *k*_*ard*_) that encode feature similarity differently. The linear combination of kernel matrices 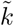 here embeds all subject-specific input data (e.g. voxel-based images) in a high dimensional space to measure similarity between each input. One key advantage of using above kernel mappings for GPR and GPC is to replace the large covariance matrices representing the high dimensional images by much smaller (and more efficient) kernel matrices (*n* × *n*).

### 2.5. Model optimization and generalization performance

The model optimization for all GP models was performed by maximizing the log-marginal likelihood using Newton-conjugate gradient descent (Bishop, 2006), which aims to find a global optimal value of hyperparameters. In order to evaluate the out-of-sample predictive performance of the GP model, we employed 10-fold nested cross-validation (CV) strategy on the whole dataset, where the outer 10-fold loop assesses the performance of the model. This strategy was followed to study the influence of feature combinations, image parameters (resolution) and kernels under various features and model combinations. The hyper-parameters of the model are evaluated using the inner 10 folds and the final CV performance of the model was measured as an average score (Pearson’s correlation and *R*^2^ score in case of regression and AUC score in case of classification) across the outer loop of 10-fold nested CV (Scheinost et al., 2019; Varoquaux, 2018; Kohavi et al., 1995; Pereira et al., 2009). We also perform training repeatedly with different initialization of parameters *θ* for finding the global minimum. All applications using GP inference and prediction on MRI data in this paper were performed using custom-made implementation in Python. The code for the GP-MKL model is available at https://github.com/neuroprognosis/GPMKL.

### 2.6. Application to real MRI sample

The DZNE-longitudinal cognitive impairment and dementia (DELCODE) cohort is a multi-centric observational study collected at 10 sites of the German Center of Neurodegenerative Diseases (DZNE). The full sample at baseline includes 1079 participants representing a broad spectrum ranging from healthy towards clinically diagnosed as dementia. More specifically these include 236 healthy controls (HC) without any cognitive impairment, 444 subjects with subjective cognitive decline (SCD), 191 cases with mild cognitive impairment (MCI), 126 Alzheimer’s patients (AD) and 82 first-degree relatives of AD patients (ADR) (Jessen et al., 2018). Subjects in the HC and ADR groups were recruited by public advertisement, whereas SCD, MCI and AD subjects through referrals (including self-referrals) in the participating memory centers. Out of 1079 subjects, 973 subjects were between 59 and 89 years old and had T1-weighted data. Of these 973 subjects, 14 subjects were excluded due to MR artifacts and poor processing quality (see details below). The inclusion criteria for the DELCODE study were as follows: SCD, MCI and AD participants were clinically assessed including medical history, psychiatric and neurological examination, neuropsychological testing, blood laboratory work-up, and routine MRI, all according to the local standards. SCD subjects were defined by the presence of subjective cognitive decline with a test performance better than -1.5 standard deviations (SD) below the age, sex, and education-adjusted normal performance on all subtests of the Consortium to Establish a Registry for Alzheimer’s Disease (CERAD) neuropsychological battery. The MCI group included individuals with amnestic MCI, defined by observed cognitive decline and age, sex, and education-adjusted performance below –1.5 SD on the delayed recall trial of the CERAD word-list episodic memory tests (Jessen et al., 2014; Molinuevo et al., 2017). Moreover, subjects with mild Alzheimer’s dementia had a score of at least ≥ 18 on the Mini-Mental-State Examination (MMSE) according to the recommendation from the National Institute on Aging-Alzheimer’s Association workgroups on diagnostic guidelines for Alzheimer’s disease (McKhann et al. (2011)).

Via advertisement text the HC and ADR participants were explicitly inquired to feel healthy and without relevant cognitive problems. All subjects that responded to the advertisement were screened by telephone with regard to SCD criteria (above). The report of very subtle cognitive decline, which (A) did not cause any subjective concerns; and (B) was considered normal for the age by the individual, was not an exclusion criterion for the HC group. Both the HC and SCD groups had to achieve unimpaired cognitive performance according to the same definition. Additional inclusion criteria for all groups were an age of at least 60 years, fluent German language skills, capacity to provide informed consent, and presence of a study partner. The descriptive statistics of 959 subjects are summarized in Table 1.

**Table 1:** Demographic information for the participants from the DELCODE cohort used in this modelling study. The memory score (see section2.7) is transformed using min-max normalization to the unit interval. Age & education are indicated in years.

| Variable | HC | SCD | MCI | AD | ADR |
| --- | --- | --- | --- | --- | --- |
| No. of subjects | 229 | 388 | 158 | 109 | 75 |
| Males/females | 129/94 | 173/200 | 69/82 | 63/44 | 44/31 |
| Age (Mean ± SD) | 69.46 ± 5.42 | 71.27 ± 6.06 | 72.91 ± 5.72 | 75.19 ± 6.25 | 66.27 ± 4.61 |
| Memory score (Mean ± SD) | 0.77 ± 0.09 | 0.72 ± 0.11 | 0.49 ± 0.14 | 0.25 ± 0.10 | 0.76 ± 0.11 |
| Education (Mean ± SD) | 14.72 ± 2.75 | 14.82 ± 2.99 | 14.11 ± 3.21 | 12.84 ± 3.04 | 14.60 ± 2.77 |

### 2.7. Memory performance assessment

The DELCODE neuropsychological battery assessment (DELCODE-NP) includes MMSE (Folstein et al., 1975), ADAS-Cog 13 (Mohs et al., 1997), FCSRT-IR (Grober et al., 2009), WMS-R Logical Memory Story A, WMS-R Digit Span (Petermann and Lepach, 2012), semantic fluency (animals) (Lezak et al., 2004), the oral form of the Symbol–Digit–Modalities Test (Thalmann et al., 2000), Boston Naming Test (Smith, 1982), Trail Making Test A and B (Reitan, 1958), Clock Drawing, and Clock Copying (Rouleau et al., 1992). In addition, the Face Name Associative Recognition Test (Polcher et al., 2017), and a Flanker task were used to assess executive control of attention (Van Dam et al., 2013). These tests were selected to have similar compatibility with ongoing studies such as ADNI (Park et al., 2012) and WRAP (Dowling et al., 2010) and to derive cognitive domain scores (includes learning and memory, language ability, executive functions and mental processing speed, working memory and visuospatial abilities) and cognitive composite score (e.g., the “Preclinical Alzheimer cognitive composite” (PACC5) (Papp et al., 2017)) for tracking cognitive decline. We calculated the preclinical Alzheimer’s cognitive composite (PACC-5) as the mean of an individual’s z-standardized performance (based on the cognitively unimpaired individuals) in the FCSRT Free Recall and Total Recall, the MMSE, the Wechsler Memory Scale-R (WMS-R) Logical Memory Story A Delayed Recall, the Symbol-Digit-Modalities Test, and the sum of the two category fluency tasks (animals, grocery). Factor scores or composites have been shown to improve predictive performance (Dubois et al., 2018).

In our study, global memory performance factor with improved psychometric properties was derived using Confirmatory Factor Analysis (CFA) for baseline measures (Wolfsgruber et al., 2020). Due to availability constraints, cross-sectional baseline predictions of memory performance were based on this CFA-based memory factor while longitudinal analyses were focussed on PACC5 score.

### 2.8. CSF biomarker positivity

Out of 959 subjects of the DELCODE cohort, Cerebrospinal fluid (CSF) biomarker (*Aβ*42/40 and pTau) characterization was available for a subset of 453 (47.23 %) subjects. The CSF biomarkers were determined using commercially available kits according to vendor specifications: V-PLEX *Aβ* Peptide Panel 1 (6E10) Kit (K15200E) and V-PLEX Human Total Tau Kit (K151LAE) (Mesoscale Diagnostics LLC, Rockville, USA), and Innotest Phospho-Tau(181P) (81581; Fujirebio Germany GmbH, Hannover, Germany). The cutoff value of the normal and abnormal concentration of *Aβ*42/40 is defined as 0.08 pg/ml (below pathological) and of pTau 73.65 pg/ml (above pathological). These cutoffs were determined on the basis of all DELCODE Baseline CSF data (n = 527) by Gaussian mixture modeling using the R package flexmix (version 2.3-15).

### 2.9. MRI acquisition

MRI scans were acquired in 9 out of 10 involved DZNE sites (3T Siemens scanners: 3 TIM Trio systems, 4 Verio systems, 1 Skyra and 1 Prisma system). Our main analyses were based on whole-brain T1-weighted MPRAGE (3D GRAPPA PAT 2, 1 mm3 isotropic, 256 × 256 px, 192 slices, sagittal, 5 min, TR 2500 ms, TE 4.33 ms, TI 110 ms, FA 7°). Further ROI and covariate processing was based on the additionally available FLAIR protocol (for details see (Jessen et al., 2018)).

### 2.10. MRI preprocessing and feature generation

We prepared morphometric brain features following ideas from computational anatomy and Voxel-based Morphometry (VBM). This was achieved by combining spatial normalization of SPM (Welcome Trust Centre for Human Neuroimaging, London, UK, http://www.fil.ion.ucl.ac.uk/spm) and segmentation from CAT12 (http://www.neuro.uni-jena.de/cat/, r1615, Jena, Germany) toolbox. All T1-weighted images were corrected for bias-field inhomogeneities, non-brain tissue was stripped, the grey matter (GM), white matter (WM), and cerebrospinal fluid (CSF) brain tissue types were segmented using the CAT12 segmentation algorithm with partial volume estimation to account for mixed voxels with two tissue types (Tohka et al., 2004) and adaptive maximum a posteriori (AMAP) (Rajapakse et al., 1997). Finally, all scans were iteratively registered with a study-specific template space using rigid and non-linear diffeomorphic transformations (Ashburner and Friston, 2009, 2011). Previous results from Monté-Rubio et al. (2018) show that unmodulated segment images when compared to modulated segment images (using Jacobian determinant) showed improved predictive performance over multiple tasks. We therefore generated and compared different morphometric features under variation of certain imaging parameters such as resolution (see next section). All further kernel estimates were based on vectorized versions of these images and represent a whole-brain approach.

### 2.11. Accounting for covariates, risk factors and nuisance variables

Covariates and nuisance variables that correlate with the imaging data might affect the predictive performance of a model by adding variability to the data (Scheinost et al., 2019; Sanderman et al., 2006). From a clinical perspective, these variables can also affect the interpretability of the model trained using neuroimaging data. Therefore, it’s important to account for these variables in the predictive model. Rao et al. (2017) suggested in dealing with confounds with respect to predictive modelling, is to include confounds as a predictor along with the imaging features in the model (Rao et al., 2015). The most commonly found demographic covariates in prediction studies are age and sex (Alfaro-Almagro et al., 2021; Rao et al., 2017; Abdulkadir et al., 2014). Importantly, the latter variables have been associated with the risk of progression towards pathological ageing and AD (Lindsay et al., 2002; Riedel et al., 2016). In addition, genetic information such as ApoE4 status has been shown to increase the risk for AD and therefore might covary with memory performance and biomarker status (Wolfsgruber et al., 2014). An important nuisance variable in multi-center studies is is the acquisition site. Therefore, in this predictive modelling study we considered acquisition site as predictors to account for potential covariate and confounding effects in the model by including as a predictor along with different types.

### 2.12. Application of GP-MKL framework to the DELCODE cohort

Next, we evaluated the performance of the GP-MKL framework for providing clinically relevant predictions of (Task I) memory performance and (Task II) biomarker positivity (*Aβ*42/40 and pTau) in DELCODE study participants. A schematic overview of the applied framework is provided in Fig. 1 following the preprocessing pipeline described earlier.

**Figure 1:**
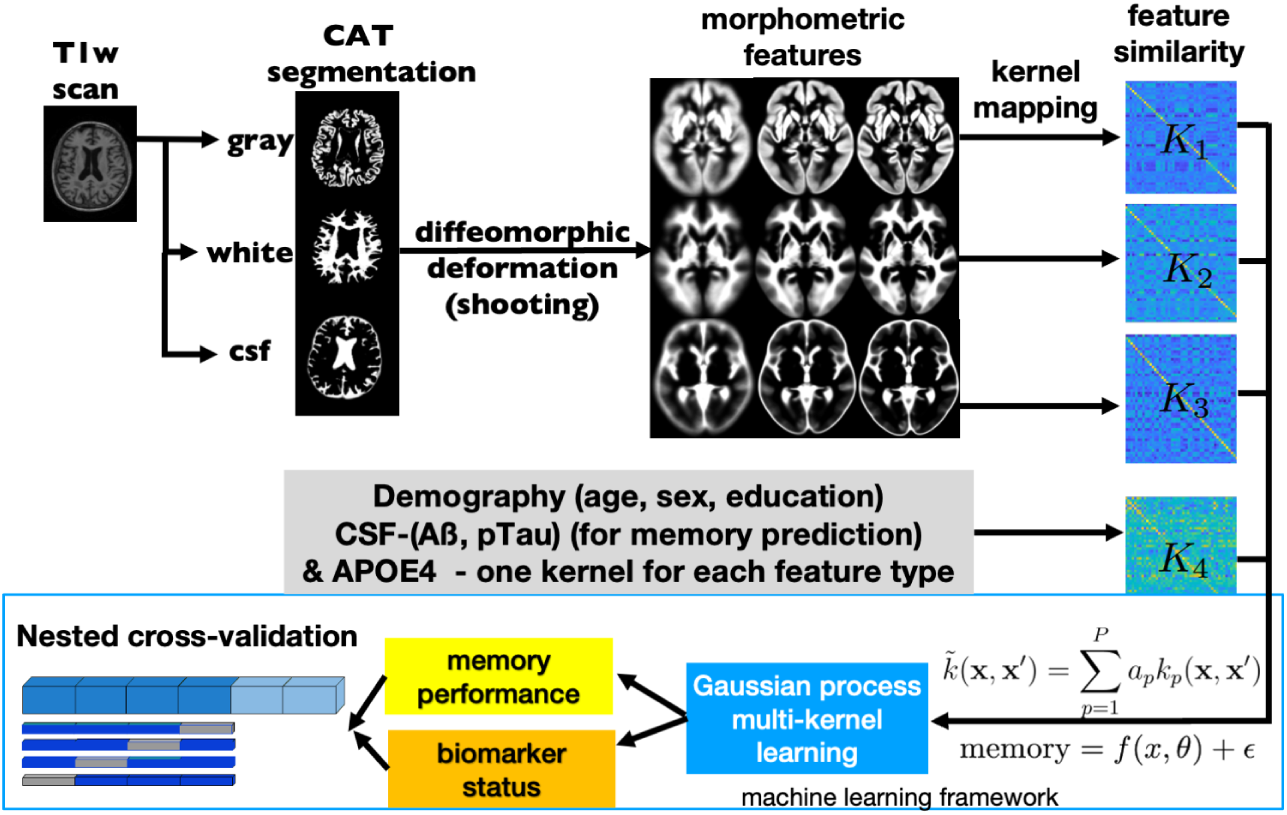
GP-MKL framework, where the T1-weighted scans are segmented into different brain tissue types using the CAT12 segmentation algorithm. (2) The segmented tissue types are then aligned in the same space using a non-linear image registration algorithm to obtain morphometric features. (3) These morphometric features are mapped to kernels and forwarded to the GP-MKL framework for prediction of memory performance and biomarker status using 10 fold nested CV.

#### 2.12.1. Prediction Task I: Memory performance

The individual memory performance was predicted using successively richer feature sets comprising (A1) demographic covariates and ApoE4 genotype (A2) multiple morphometric imaging features such as gray matter density; and (A3) CSF biomarkers. First, we started by using commonly known informative demographic covariates (Demo = [age, sex, education]) and derived one separate kernel for each of these feature types entering the GP-MKL model. Additionally, we include ApoE4 a strongest genetic risk factor of AD (M Di Battista et al., 2016). This forms a baseline model which captures those cognitive performance differences.

Second, we focused on memory prediction using MRI-based brain features. More specifically, we systematically compared (A) combinations of tissue classes (GM, WM, CSF) and feature type (modulated vs. unmodulated) for memory performance prediction; (B) the influence of smoothing kernel size; and (C) image resolution (1mm, 2mm, 4mm and 8mm) for the resulting brain tissue features fed into GP-based predictive models. Initially, we applied a wholebrain mask to extract voxelwise features such as gray matter density for each subject. The features were then fed into selected kernels (eq. 4, 5, 6) for different feature types to derive kernel matrices. These kernel matrices were then used to predict memory performance and biomarker positivity using GP-MKL.

Third, since neuropsychological testing, MRI and CSF biomarkers come at different costs for patient and researchers, we aimed at quantitative head-to-head comparisons. For the subsample with available biomarker characterization, we additionally generated kernels for *Aβ*42/40 and pTau separately using eq. 4, 5, 6. Note, the biomarker-based predictive features are only used during our first task aiming at memory performance as a target variable. The above potential sources of information are used to predict performance differences are expected to be partially redundant or complementary. We therefore, evaluated their selected feature combinations, resulting in more complex models and a higher number of kernels. In order to assess the performance and validate the GP-MKL predictive model during nested 10-fold CV, the correlation between predicted and true memory score and prediction *R*^2^ (cross-validation *R*^2^ = 1 −*MSE*(*predicted* −*observed*)*/MSE*(*observed* −*μ*)) score was estimated, where MSE is the mean squared error and *μ*is the mean of the observed variable (Scheinost et al., 2019).

#### 2.12.2. Prediction Task II: Biomarker positivity

To classify biomarker positivity (*Aβ*42/40(+ve/-ve) and pTau(+ve/-ve)) for the available data of 453 subjects, we used the same combinations of demographics, ApoE4 genotype and MRI features that we used during prediction of memory performance. To avoid circular inference, CSF biomarker features were not used as inputs to the machine learning algorithm in this prediction task. However, we additionally included the memory factor score obtained from neuropsychological testing to the set of evaluated feature combinations. For prediction of binary biomarker positivity, the overall performance was assessed by determining how well the classifier performed at various threshold settings. This was done by evaluating the area under the curve (AUC) of the Receiver Operating Characteristic (ROC). If AUC=1, the classifier can distinguish +ve and -ve classes in all test subjects perfectly. Since the classifier operates at 50-50 chance of guessing the classes correctly, if 0.5 *< AUC <* 1, there is a better chance at identifying classes accurately than guessing. Note, before evaluating the performance of the classifier, the training and test samples were balanced across the outer and inner fold of the nested cross-validation.

#### 2.12.3. Testing GP-MKL model variations

The above tasks were focussed on evaluating predictive combinations of input feature sets. However, there are potential contributions of preprocessing parameters. Firstly, we studied choices of filter size of smoothing (ranging from 0 to 15 mm FWHM) using modulated and unmodulated brain tissue types for various image resolutions (1mm, 2mm, 4mm and 8mm). Additionally, we studied the influence of further aspects of the model and training sample. We investigated the impact of the size of the training data set (i.e., number of subjects) in order to enable useful extrapolations for future studies on similar prediction tasks. Moreover, as shown in previous machine learning applications we explored the potential influence of reducing the dimensions of data using principal component analysis (PCA). Finally, we test the particular choice of linear vs. non-linear kernels such as squared exponential kernel (aka RBF) and ARD kernel.

#### 2.12.4. Validation using longitudinal data

The above model explored baseline associations of brain and clinical outcomes using a predictive approach. However, the framework has potential for clinical translational applications by predicting clinical outcome measures not only at baseline but at follow up measurements, i.e. prediction of the future memory scores. In this final experiment, we used available longitudinal data of the DELCODE cohort to predict individual memory performance at follow-up using baseline MRI and other covariates. Due to longitudinal availability, we here focussed on the cognitive composite score PACC5 (rather than the memory factor score) and re-trained the model using baseline data (DEMO, MRI and CSF biomarkers) to predict PACC5 scores at annual followups. The PACC5 score was available at baseline and five annual follow-ups for 877/695/502/373/197/41 subjects respectively.

## 3. Results

### 3.1. Prediction of memory performance

We started by training the GP-MKL framework with a simple baseline model that only used informative demographic covariates (DEMO) such as age, sex and education to predict each participant’s individual memory performance. The model performance in terms of correlation of predicted and true memory scores and predictive *R*^2^ averaged across outer 10-fold CV was estimated to be *r* = 0.49 ± 0.11 and *R*^2^ = 0.24± 0.1. Additionally including ApoE4 improved the performance of the model to *r* = 0.53 ± 0.09, *R*^2^ = 0.28 ± 0.09. Therefore, we further included ApoE4 as one of the demographic covariates (abbreviated as DEMO+ApoE) in all further analyses.

We then explored the potential of using MRI-based brain features for improved prediction of memory performance over the above baseline model. There is a variety of morphometric voxel-based MRI features such as tissue class, modulation vs. no-modulation, smoothness etc. that can be chosen for MKL predictive models. We studied the effects of variations of pre-processing of imaging features in greater detail including the results of optimal parameters presented in a subsequent section (3.3) and here focus on the summary how demographic covariates and risk factors in combination with these ‘optimal’ sMRI imaging features (GM+CSF of 4mm smoothness and image resolution) performed in predicting the participant’s memory performance. A summary of out of sample predictive performance for all combinations of feature types in predicting memory is provided in Table 2.

**Table 2:** Illustrates summary of predictive accuracy for different combination of feature types to predict individual memory performance. The predictive variance refers to the variance of the predictive distribution given the baseline features of an unseen subject is given as a measure of uncertainty of individual predictions. When CSF biomarkers were used it also increases since training sample size is reduced due to availability (see also Fig 3.3 (b.1) for the effect of predictive variance for different sample sizes). Kernels (hyperparameters) represents no: of kernels used in the model and no: of hyperparameters estimated by the model.

| Feature types | Performance scores |  |  | Kernels (hyperparameters) |
| --- | --- | --- | --- | --- |
| | Correlation score | $R^2$ score | Predictive variance | |
| DEMO+ApoE4 | $0.53 \pm 0.09$ | $0.28 \pm 0.09$ | 0.1042 | 4(4) |
| sMRI | $0.73 \pm 0.03$ | $0.51 \pm 0.05$ | 0.0755 | 2(2) |
| CSF Biomarkers | $0.56 \pm 0.13$ | $0.32 \pm 0.14$ | 0.2206 | 2(2) |
| DEMO + ApoE4 + sMRI | $0.75 \pm 0.02$ | $0.55 \pm 0.04$ | 0.0754 | 6(6) |
| DEMO + ApoE4 + CSF Biomarkers | $0.64 \pm 0.08$ | $0.40 \pm 0.1$ | 0.2203 | 6(6) |
| sMRI + CSF Biomarkers | $0.75 \pm 0.07$ | $0.55 \pm 0.1$ | 0.1564 | 4(4) |
| DEMO + ApoE4 + sMRI + CSF Biomarkers | $0.77 \pm 0.06$ | $0.57 \pm 0.1$ | 0.15645 | 8(8) |

The combination of demographics, ApoE4 and optimal sMRI revealed a predictive accuracy of *r* = 0.75 ± 0.02 and predictive *R*^2^ = 0.55 ± 0.04. Therefore, including specific morphometric sMRI features in addition to demographic variables enhanced the overall performance substantially by Δ*R*^2^ = 0.27. However, adding subject level risk scores like age, sex, education and ApoE4 did only slightly improve predictive performance of memory performance over sMRI in isolation.

Finally, we incorporated the CSF biomarker (47.23% subjects) characterization in our predictive framework for the memory prediction task. More specifically, we focused on the question if *Aβ*42/40 and pTau levels obtained from invasive lumbar puncture along with demographic and/or sMRI can further improve prediction of each participant’s memory performance. For this purpose, we evaluated the GP-MKL model in a sub-sample with available CSF biomarkers and generated separate kernels for *Aβ*42*/*40 and pTau. When using demography, ApoE4 status, sMRI and CSF biomarkers combined nested cross validation revealed accuracies of *r* = 0.77 ± 0.06 and predictive *R*^2^ = 0.57 ± 0.1. The relationship between predicted and true memory performance for this model is illustrated in Fig. 2.

**Figure 2:**
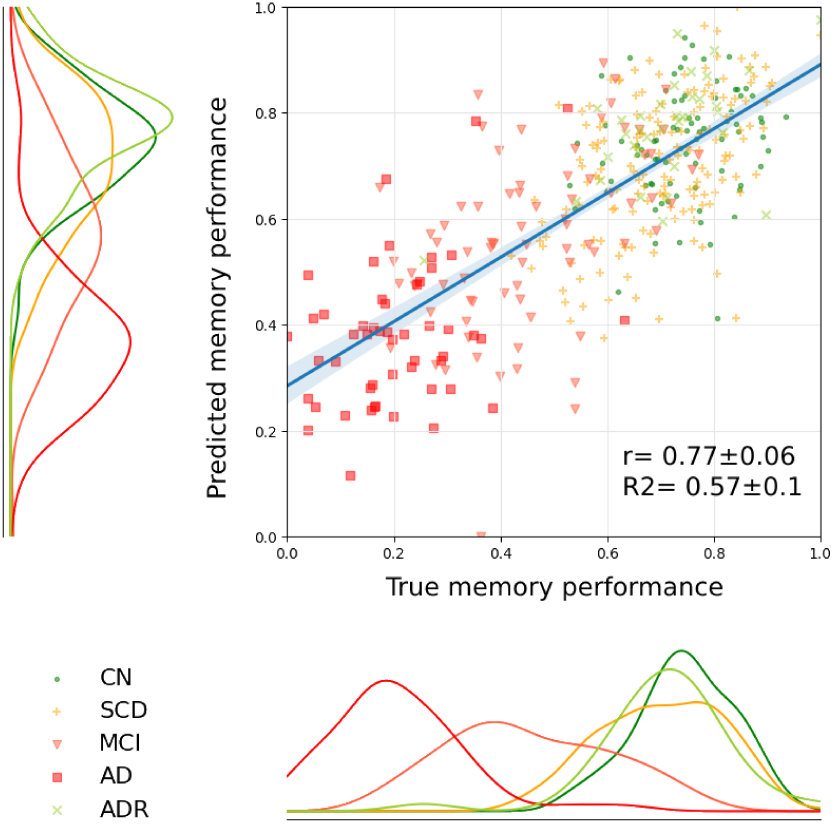
Illustrates prediction of individual memory performance using a combination of demographic covariates, MRI-based brain features (4mm image resolution and 4mm FWHM smoothness), and CSF biomarkers (*Aβ*42*/*40 and pTau).

In summary, the overall predictive accuracy improvement over the baseline model, which only considered covariates (e.g. from demography and ApoE4 status) increased by Δ*R*^2^ = 0.27 for MRI and Δ*R*^2^ = 0.12 for inclusion of *Aβ*42*/*40 and pTau biomarkers. The combination of demographic, structural brain and CSF biomarkers feature types revealed highest predictive accuracies in predicting the baseline memory performance. Furthermore, the variance of the predictive distribution was slightly improved for the best predictor combinations (see Table 2).

### 3.2. Prediction of biomarker positivity

In analogy to the above procedure, we first used the informative covariates (such as age, sex, education, and ApoE4) to predict CSF biomarker positivity. We trained our GP-MKL classifier for the DELCODE cohort subsample with available CSF biomarkers (42.07% of subjects) and assessed the classifier performance using nested cross-validation.

The accuracy for prediction of *Aβ*42*/*40 positivity when only using these covariates was estimated with an AUC score of 0.83 ± 0.06 (specificity: 75%, sensitivity: 77%) and for pTau an AUC score of 0.73 ± 0.07 (spec: 62%, sens: 75%). Similar to the memory prediction task, we explored including variations of voxel-based MRI features with different tissue classes, modulation vs no-modulation and various smoothness parameters (for explanation see section 3.3). We then used the best performing imaging features from the biomarker classification task in addition to covariates to enrich the inputs.

Using the combined features from sMRI, demographics, and ApoE4, we observed a classifier performance with AUC=0.76 ± 0.12 (spec: 76%, sens. 61%) for *Aβ*42/40 and AUC=0.82 ± 0.13 (spec: 94%, sens: 38%) for pTau. The overall accuracy of pTau classification was slightly improved when using additional sMRI imaging features while accuracy for *Aβ*42*/*40 was unexpectedly found to be reduced. The increase of AUC for pTau was accompanied by a marked increase in specificity to 94% while the sensitivity dropped to 38%. To investigate what’s potentially driving the sensitivity differences, we trained the classification model on imaging data alone and achieved an AUC score of 0.72 ± 0.12 (spec: 75%, sens: 54%) for *Aβ*42*/*40 and AUC score of 0.80 ± 0.13 (spec = 94%, sens = 34%) for pTau. The results suggested that the pattern of very high specificity and low sensitivity is rather a characteristic of the structural sMRI features during pTau classification. The AUC values for selected feature combinations are provided in Table 3.

**Table 3:** Illustrates a summary of Area Under Curve (AUC) performance scores from nested cross-validation for different combinations of features to classify CSF biomarkers positivity using the GP-MKL framework in the DELCODE cohort.

| Feature types | Performance scores |  |
| --- | --- | --- |
| | AUC score $A\beta_{42}/40$ (spec/sens) | AUC score pTau (spec/sens) |
| *DEMO + ApoE4* | $0.83 \pm 0.06$ (75%/77%) | $0.73 \pm 0.07$ (62%/75%) |
| *sMRI* | $0.72 \pm 0.12$ (75%/54%) | $0.80 \pm 0.13$ (94%/34%) |
| *Memory score* | $0.76 \pm 0.1$ (78%/61%) | $0.76 \pm 0.1$ (71%/71%) |
| *DEMO + ApoE4 + sMRI* | $0.79 \pm 0.12$ (81%/63%) | $0.82 \pm 0.13$ (94%/38%) |
| *DEMO + ApoE4 + Memory* | $0.86 \pm 0.06$ (81%/76%) | $0.78 \pm 0.07$ (70%/74%) |
| *DEMO + ApoE4 + Memory + sMRI* | $0.78 \pm 0.12$ (79%/63%) | $0.83 \pm 0.12$ (95%/37%) |

To further explore alternative feature combinations for optimal CSF biomarkers classification, we used neuropsychological test-based memory performance scores to predict CSF biomarker positivity. Notably, while playing the role of the target in above prediction task, here memory performance is an additional input to classification algorithms predicting biomarker status. We combined kernels for memory performance with those from demographic and ApoE4 covariates (as above) and obtained an estimate of classification accuracy for *Aβ*42/40 of AUC=0.86 ± 0.06 (spec: 81%, sens: 76%) and for pTau the AUC=0.78 ± 0.07 (spec: 70%, sens: 74%). Using a neuropsychological memory score provided higher predictive accuracy (specificity and sensitivity) when classifying CSF *Aβ*42*/*40 positivity compared to using neuroimaging markers. For pTau the integrated AUC of 0.82 was slightly higher when using sMRI instead of AUC of 0.78 when using a memory test-score (in combination with demographic covariates). However, the sensitivity and specificity profile was more un-balanced when using sMRI pointing to very high specificity at the cost of sensitivity.

Finally, we combined all available baseline feature sets using neuropsychological memory performance, with demographic covariates and morphometric brain features. The classifier performance was estimated as AUC=0.78 ± 0.12 (spec: 79%, sens: 63%) for *Aβ*42/40 and AUC=0.83 ± 0.12 (specific: 95%, sens: 37%) for pTau. The combination of all features only improved performance in case of classifying pTau while accuracy for *Aβ*42*/*40 positivity was found to be reduced compared to the above reported of combination of covariates with a neuropsychological-based memory score without MRI.

### 3.3. Study variations of MRI features, sample size, kernel type, and PCA

Since feature engineering can be essential when constructing powerful ML models (Franke et al., 2010; Monté-Rubio et al., 2018) we also studied choices of (A) filter size of smoothing (ranging from 0 to 15 mm FWHM); (B) using modulated or unmodulated brain tissue segments; (C) which selection of tissue class(es) among GM, WM, and CSF; and (D) various image resolutions (1mm, 2mm, 4mm and 8mm). Based on smoothed images of varying filter size we derived separate linear kernels for each feature type and brain tissue class which were subsequently entered as a linear combination in the predictive modelling framework.

The effects of various combinations of input features and parameters on accuracy of MRI-based memory prediction (i.e. only using MRI-derived features without (demographic) covariates or CSF) is summarized in Fig. 3a.

**Figure 3:**
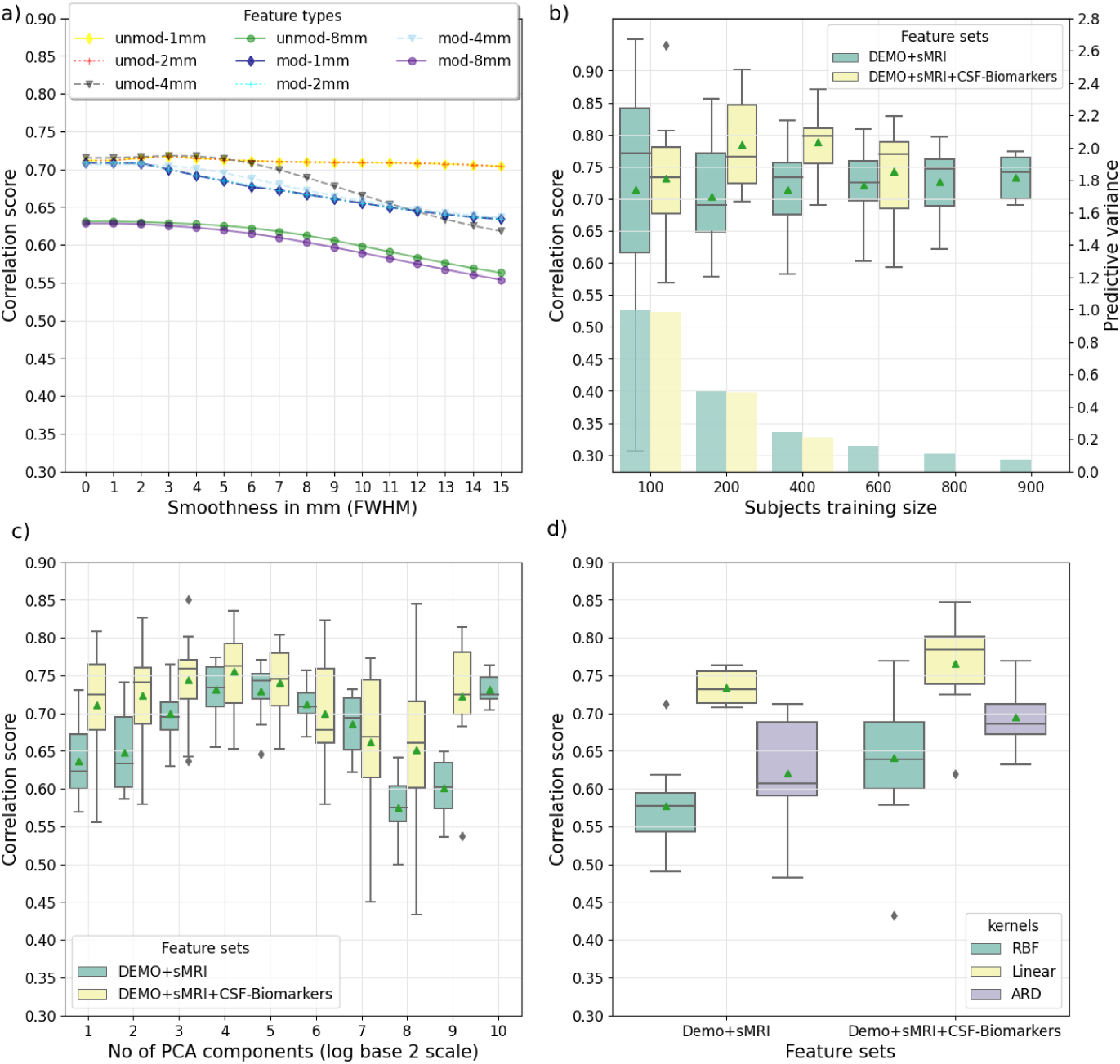
Illustration of effects of choice of morphological features, training sample size, PCA and kernel type on memory performance prediction using GP-MKL framework (a) Evaluating the influence of smoothing kernel, image resolution, combined tissue classes (GM+WM+CSF) and feature type (modulated vs. unmodulated). Only MRI ; (b) Impact of training sample size; (b-yyaxis) Effect of predictive variance w.r.t sample size; (c) Influence of reducing the dimensionality of data using Principal Component Analysis (PCA), numbers of components in log_2_ scale; and (d) Evaluating the predictive accuracy when using linear, squared exponential kernel (aka RBF) and ARD kernel.

Our results indicated that the combination of GM and CSF brain tissue classes for unmodulated segments at 4 mm image resolution and 4 mm smoothness (FWHM) performed best for the memory prediction task (see also previous Table 2). For CSF-biomarkers classification, combination of GM and CSF unmodulated tissue types performed best at 2 mm image resolution with 8 mm smoothness for *Aβ*42*/*40 and 6 mm smoothness for pTau prediction. It is worth noting that the smoothing filter size, feature types (modulated and unmodulated) and selection of tissue classes might perform differently for other predictive tasks.

MR-based predictions using unmodulated tissue segments revealed generally higher accuracies when compared to modulated segments across various smoothing kernels and image resolutions across both the predictive tasks. The predictive accuracy for predicting memory performance, when using 1 mm and 2 mm image resolutions was found to be comparable for most filter sizes, but slightly lower compared to 4 mm with smaller smoothing kernels. Predictive accuracy remained comparably unchanged for various smaller smoothing kernels but declined gradually with more smoothing. The 8 mm input resolution generally revealed worst overall results.

In addition, we separately evaluated different combinations using GM+WM+ CSF, GM+WM, WM+CSF and GM+CSF. The predictive accuracy for memory prediction, when combining GM+CSF (*r* = 0.73 ± 0.03, *R*^2^ = 0.51 ± 0.05) was found to be slightly above the full combination of all tissue classes, ie. GM+WM+CSF (fixing all other manipulated parameters to optimal values described above). For CSF biomarkers classification, the classification accuracy revealed the same set of tissue types combination (for *Aβ*42*/*40-GM+WM+CSF: AUC=0.70 ± 0.12, GM+CSF: AUC=0.72 ± 0.12; pTau-GM+WM+CSF: AUC= 0.79 ± 0.13, GM+CSF: AUC=0.80 ± 0.13). Consequently, we excluded the WM tissue class in further sMRI analyses.

We further evaluated the influence of nuisance variables by including scanning site as an additional predictor to the baseline feature set and observed a memory prediction accuracy of *r* = 0.53 ± 0.09, *R*^2^ = 0.28± 0.09. The predictive accuracy therefore remained unchanged, when confounding effects of site were incorporated as inputs to the GP-MKL models for memory prediction.

We further studied the potential influence of the available numbers of observations during training on predictive outcomes. We re-trained the GP-MKL model on various training sample sizes ranging from 100 to 900 subjects using MRI combined with other feature sets (fixing other parameters on described optimal values) (Fig. 3(b)). Our findings suggested with exception of the smallest sample that performance slightly increased with larger sample and estimates appeared to converge on consistent estimates when using at least 500 subjects. As expected, the nested cross-validation accuracy estimates showed less variation across folds when using larger sample sizes compared to smaller ones. Additionally, we explored the variance of predictive distribution (see Fig 3 (b.1)) for different sample sizes for the best obtained feature combinations from the memory prediction task. Our result indicate, the variance (smaller the better) improves with larger samples. Next, we explored the potential benefits of reducing the dimensionality of the imaging data by using PCA before calculating kernels. We started by creating principal components that are uncorrelated dimensions with maximum variance and then fit the model for the best combination of image features from the memory performance task and achieved an average performance score of *r* = 0.64 ± 0.05, *R*^2^ = 0.4 ± 0.07. Similarly, we created PCA components until subjects *<*= training data. The results in Fig. 3(c) shows that when an optimal number of PCA components is chosen a-priori or selected via cross-validation, in our case the best performance was achieved for PCA components = 32 (*log*_2_(4)) with an average performance score of *r* = 0.74± 0.04, *R*^2^ = 0.50 ± 0.1. Altogether, the estimated predictive accuracy was lower than that of a non-PCA version (no dimensionality reduction) by Δ*r* = 0.03, Δ*R*^2^ = 0.07 (see table 2). Finally, we evaluated our model performance using non-linear RBF and ARD kernels and compared it with linear kernels (Fig. 3(d)). Findings indicate that linear kernels outperform both RBF and ARD kernels and ARD performs better than RBF kernels. Thus in this predictive modelling of memory task the linear kernel seems to optimally capture the input feature patterns compared to commonly used alternative kernels.

Linear kernels do allow visualizing weights corresponding to the contribution of each voxel during estimating predictive performance (see Appendix A for more details). Fig. 4(A) shows the GP-MKL weight map for prediction of memory scores using the best sMRI features obtained from the regression task.

**Figure 4:**
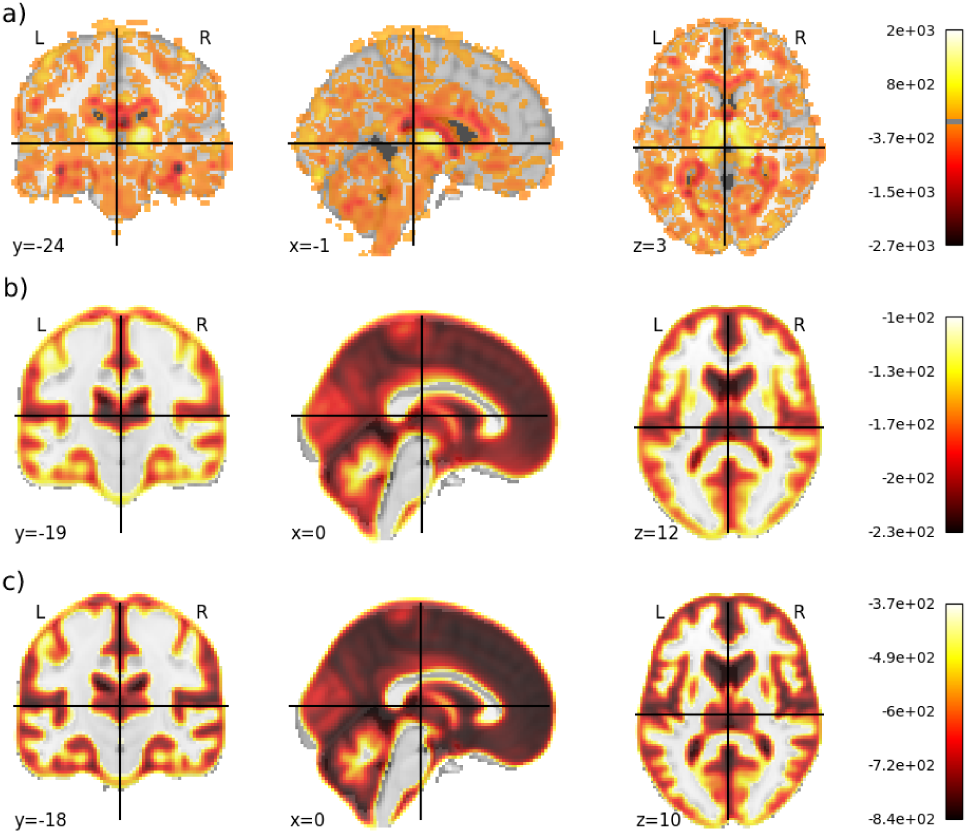
(A) Illustrates the spatial weight map from sMRI features contributing to the prediction of memory performance. (B) The weight map for sMRI features during classification of *Aβ*42*/*40 positivity; and (C) classifying pTau positivity. Higher value on the color bar represent higher weights contribution.

The results show higher weight contribution in the brain regions of thalamus, hippocampus, right fusiform gyrus and middle temporal gyrus. The corresponding weights for the classification of CSF biomarkers for *Aβ*42*/*40 and pTau classification is shown in Fig. 4(B) and (C) respectively. The higher weights contribution was found in the brain regions of cerebellum, caudate nucleus, mammillary bodies and hippocampus for *Aβ*42*/*40 prediction; cerebellum, frontal gyrus and hippocampus for pTau classification.

### 3.4. Validation using longitudinal data

In this final experiment, we used available longitudinal data of the DEL-CODE cohort to predict individual memory performance scores at follow-up using baseline features including MRI. Notably, the model makes predictions (over time) and is re-trained using baseline data including demographic covariates (age, sex, education and ApoE4), MR-based morphology, CSF *Aβ*42*/*40 and pTau to predict PACC5 scores at annual follow-ups (see methods).

In this prediction over varying time gaps, we obtained the highest performance of *r* = 0.71 ± 0.07, *R*^2^ = 0.48 ± 0.09, when using all baseline features and estimated performance of *r* = 0.69 ± 0.08, *R*^2^ = 0.43 ± 0.08, *r* = 0.56 ± 0.14, *R*^2^ = 0.28 ± 0.18, *r* = 0.47 ± 0.3, *R*^2^ = 0.2 ± 0.3 and *r* = 0.2 ± 0.5, *R*^2^ = −0.08 ± 0.04 for follow-ups respectively (shown in Fig. 5(left). Please note that due to availability PACC5 instead of the memory factor was used during this analysis. As expected the accuracy of prediction strongly drops over longer time intervals, especially after 2 years and more. Furthermore, we additionally included baseline neuropsychological memory score and estimated highest performance for combination of demographics, ApoE4 genotype and neuropsychological score (see Figure 5(right)).

**Figure 5:**
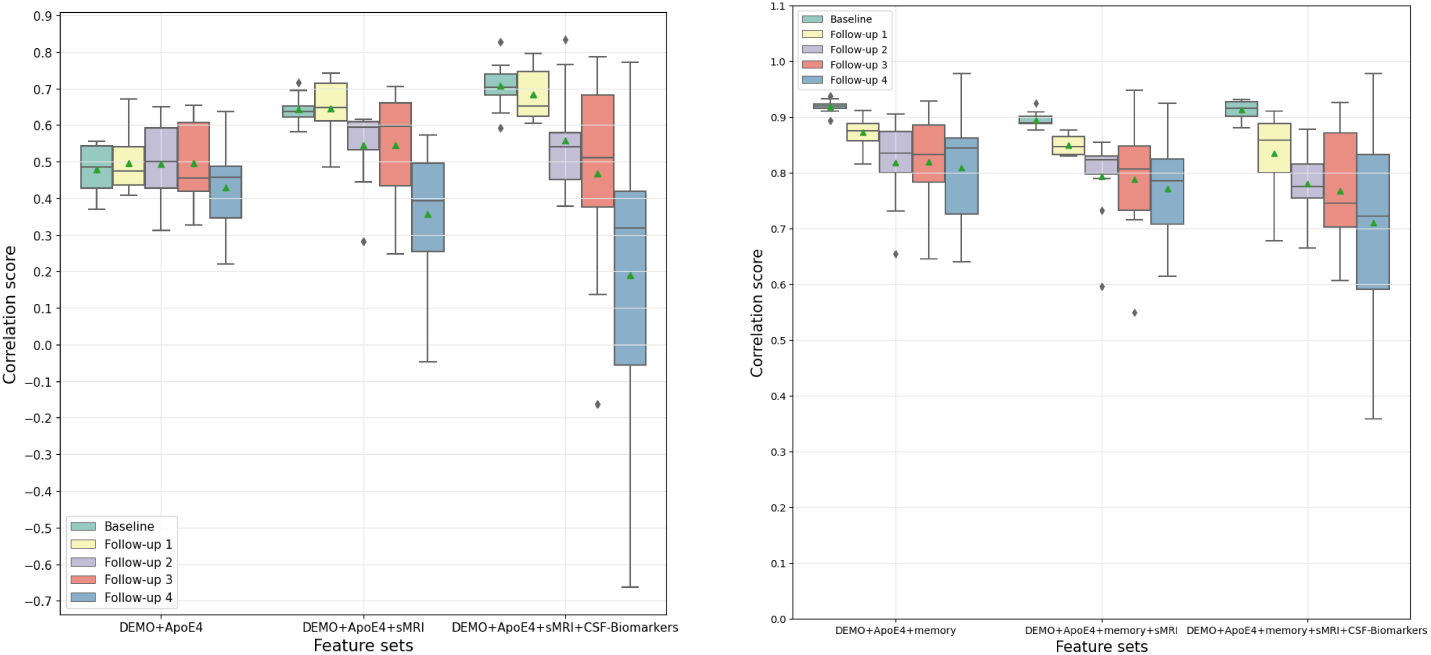
Illustration of memory performance (PACC5) prediction using DEMO+ApoE4, sMRI, CSF-biomarkers for longitudinal data (left) and additionally using baseline neuropsychological memory performance (right).

## 4. Discussion

We presented a MR-based predictive modelling of memory performance and biomarker positivity in ageing subjects at risk of Alzheimer’s disease using a large sample multi-centric observational cohort (DELCODE) and a Gaussian process multi-kernel framework. One benefit of using kernel methods is the efficiency of the approach by mapping the high dimensional brain image data (X) to the dimensions of the number of samples (n, representing number of subjects in our case), which is much lower than the actual image dimensions. Earlier studies have demonstrated the potential of kernel methods in the context of neuroimaging data (Ashburner and Klöppel, 2011; Jollans et al., 2019) especially combining multiple kernels to represent various types of predictive features information (Zhang et al., 2012; Gupta et al., 2019; Liu et al., 2018; Gönen and Alpaydın, 2011; Wilson and Adams, 2013; Monté-Rubio et al., 2018; Dyrba et al., 2015).

More specifically, to explore the beneficial combinations of information from different sources during prediction of memory performance and biomarker status we combine multiple input features using kernels for each feature type (such as brain morphometric features, subject-specific covariates and CSF biomarkers). Our investigation revealed, when using these feature types in isolation to predict memory performance, CSF-biomarkers predicted better compared to just demographic covariates and ApoE4. However, as isolated feature domains, the structural MRI (i.e. morphometric aspects of the brain) revealed the highest predictive accuracy. These results are in line with previous studies (Zhang et al., 2012; Stonnington et al., 2010; Zhang et al., 2011; Wang et al., 2010; Duchesne et al., 2009; Liu et al., 2018) that have demonstrated the use of sMRI to predict clinical cognitive scores. The proposed method, in Zhang et al. (2012), used multi-modal data that includes features from MRI, PET and CSF to achieve a correlation score of 0.697 ± 0.022 and 0.739 ± 0.012 in estimating MMSE & ADAS-Cog scores respectively. However, our method using sMRI independently achieved similar performance with a correlation score of 0.73 ± 0.03 (see Table 2) in estimating cognitive score.

A first question we aimed at was MR-based prediction of memory performance in healthy aging subjects and subjects at risk of AD. Although memory performance at baseline can be assessed via neuropsychological testing directly, this prediction of the cognitive domain (at the same time-point) forms the basis for highly relevant longitudinal predictions over time. Our study revealed evidence that the full combination information from sources such as demographic information (age, sex, and education), ApoE4, sMRI and CSF biomarkers (A*β*42/40 and pTau) offers the overall highest predictive accuracy concerning to cognitive memory performance at baseline. Using our GP-MKL approach and all input features jointly 57% (r=0.77 ± 0.06,*R*^2^=0.57 ± 0.1) of individual memory performance outcome variability could be correctly predicted using an assessment of the out of sample generalization performance via nested CV. In comparison to Zhang et al. (2012), proposed method (MMSE: r=0.697±0.022; ADAS-Cog: 0.739±0.012), our method achieved slightly higher performance in predicting memory. One might speculate that the overall predictive performance could have been further improved if CSF biomarkers were available in a larger sample.

Blood testing for ApoE4 status assessment, sMRI and biomarkers have different (dis-)advantages in terms of being invasive, actual financial expenditure, and time investment. A good cost-accuracy trade-off can be also achieved with 55% (r=.75) explained variance by excluding the CSF biomarkers from the predictive framework. This supports relevance of the MR-based approach and its potential extensions for translational clinical applications e.g. in memory clinics. Our additional analyses suggested that even larger training sample size might further increase performance slightly in future studies. Taking advantage of a Bayesian approach our GP-MKL model goes beyond enabling point estimates for memory performance in unseen subjects (as e.g. with most CNNs) by providing additional uncertainties of individual predictions (Salvatore et al., 2016; Korolev et al., 2016; Challis et al., 2015). The conducted analyses suggested that the uncertainty of predictions might be further decreased with increasing training sample size.

The finding that patterns of brain volumes were found to be more predictive of memory performance than Amyloid and pTau CSF-biomarkers (alone or in combination with covariates) might have several non-exclusive causes. First, CSF biomarkers might reflect sings of early brain pathology in terms of presence of certain proteins (Hu et al., 2007) which do not necessarily need to manifest in cognitive consequences. However, MRI can directly reflect the existing macrostructural tissue loss and atrophy in hippocampal networks (Eustache et al., 2016). Second, widespread atrophy patterns are high-dimensional and might be more reliably captured compared than the two-dimensional CSF biomarkers considered here. Thirdly, MRI anatomical patterns might additionally capture aspects of a brain reserve i.e. network anatomy differences that contribute positively to cognitive differences in face of brain pathology (Stern et al., 2020; Bartrés-Faz and Arenaza-Urquijo, 2011). In conclusion, the neuroimaging markers, demographics, genetic information and CSF-biomarkers were the best predictors of cognition in our sample. Therefore, sMRI might be considered for building decision support systems in clinical settings.

Next we evaluated MR-based classification of biomarker positivity using demographic covariates, ApoE4 and neuropsychological memory performance. When classifying *Aβ*42*/*40 positivity, the demographic covariates were found to be the best single predictor. Demographics combined with memory score showed the highest predictive performance which is in line with many previous reports (Tosun et al., 2016; Jansen et al., 2018; Insel et al., 2016; Buckley et al., 2019; Ko et al., 2019; Maserejian et al., 2019; Lee et al., 2018; Ba et al., 2019; Ansart et al., 2020). Compared to other studies our framework achieved a slightly higher AUC score with well-balanced specificity and sensitivity scores. Classification of *Aβ*42*/*40 based on sMRI was less accurate, which has been previously observed (Tosun et al., 2013, 2016; Ansart et al., 2020; Ezzati et al., 2020). Combining MRI with demographics and memory score resulted in slightly reduced the overall accuracy. Demographic covariates and memory test scores are easily acquired and a model with those inexpensive variables was found to performs best which yields a useful translation to the clinical setting. In contrast, our findings when predicting pTau positivity revealed the highest predictive accuracy for sMRI in isolation, with high specificity and low sensitivity indices. The high specificity (fewer false positive) combined with low sensitivity (higher false negative) might suggest that pTau becomes abnormal before neurodegeneration is severe. This is in line with the cascade model of biomarkers during progression towards AD (Jack Jr et al., 2013b). The combination of demographics, ApoE4, and the neuropsychological memory score was found to be the overall best performing classifier for pTau. In order to meet clinical demands, the threshold of the classifier to distinguish biomarkers status can be adjusted based on clinical relevance where high specificity might be preferred over high sensitivity.

Feature selection is an essential step to develop a powerful predictive model. Comparisons revealed that the segmented unmodulated GM and CSF brain tissue types in combination performed better during predicting memory and CSF biomarkers positivity compared to using modulated segments. Monté-Rubio et al. (2018) reached similar conclusions where the modulated (Jacobian-scaled) tissue types performed poorly in comparison. We observed that WM in addition to other tissue types did not provide benefits in terms of predictions which aligns with previous work Stonnington et al. (2010). The combination of smoothness filter size of 4mm FWMH and image resolution of 4mm delivered slightly better performance in predicting memory than other image resolutions and smoothness filters. However, the optimal smoothing filter size and image resolution were found to be different across tasks again suggesting importance of feature engineering in this kernel based approach. The comparisons revealed an optimal performance when using 2mm image resolution and a smoothness filter size of 8mm for classification of *Aβ*42*/*40 and 6mm for pTau. It is important to highlight the fact that the impact of predictive performance improves slightly with the use of a large training sample. Additionally, we achieved a better predictive variance (i.e. smaller uncertainty) with larger training samples and might generally minimize danger of model over-fitting. The results when comparing different kernels indicated slightly improved performance using the linear kernel in capturing input feature patterns. However, when exploring the potential of reducing dimensions of imaging data using PCA the performance was slightly lowered compared to models without PCA.

An important question associated with the interpretation (Grosenick et al., 2013) and transparency of the predictive model is how well the framework characterizes the brain regions that drive the predictive performance. The framework allows us to visualize weights that determine the association of underlying voxel signals in predicting memory performance or biomarker positivity. For memory prediction, higher weight contribution is seen in the brain regions of thalamus, hippocampus, right fusiform and middle temporal gyrus. In case of CSF biomarkers prediction higher weights are found in the brain regions of cerebellum, caudate nucleus, mammillary bodies and hippocampus for *Aβ*42*/*40 prediction; cerebellum, frontal gyrus and hippocampus for pTau classification. However, as pointed out by Mourao-Miranda et al. (2005), the weights do not assess the information in brain voxels directly and measure the corresponding contribution of each voxel in making a predictive decision. Accordingly, the relationship between the weights and neuroscientific processes should be interpreted carefully (Marquand et al., 2010).

Finally, prediction of the memory performance score (PACC5) at annual follow-ups based on baseline features to explore the associations of baseline brain and change in cognitive performance achieved potentially clinically useful estimates up to 2 years after baseline. In a previous study Wang et al. (2010) used baseline MRI to predict future decline in cognitive performance and achieved a correlation score of 0.54, Zhang et al. (2012) proposed a Multi-Modal Multi-Task (M3T) learning method to predict cognitive decline in 2 years to achieve a correlation score of 0.522. In contrast, our framework achieved slightly higher performance (r=0.54) in predicting cognitive decline in 2 years.

The study comes with few limitations, one of them is the limited availability of CSF biomarkers. Although including biomarkers data already increases the performance in predicting memory score (shown in Table 2). We believe, that availability of CSF biomarker data in the whole sample would improve the model performance even more. Furthermore inclusion of additional modalities like fMRI, quantitative susceptibility maps (QSM) and White Matter Hyperintensities (WMH) etc. could yield further improvements in predictive performance, which will be subject of future analyses which could improvise the performance of the model as shown in previous studies (Zhang et al., 2012; Gupta et al., 2019; Bouwman et al., 2007; Chételat et al., 2005). Another limitation is, only PCA was used to reduce dimensionality, in contrast other efficient dimensionality reduction techniques could be useful and reduce the complexity in data and might offer us better overall predictive performance.

In summary, our framework uses the powerful kernel methods to identify the optimal set of feature type combinations to predict baseline and follow-up memory performance, and CSF biomarker status. The framework might support reliable decision-making process e.g. for potential treatments by introducing a more precise expectation of a clinical outcomes. Future studies might explore the influence of predictive performance by including other (also longitudinal) imaging modalities and selecting feature combinations in order to further push the limits of predictions of (future) memory performance and biomarker status.

## Acknowledgements

The study was supported in part by the German Center for Neurodegenerative Diseases (DZNE), Study-ID: DZNE BN012

## Author contribution

A.N. carried out the experiment. A.N., G.Z, E.D. wrote the manuscript. G.Z, E.D, M.D., A.M., D.B., N.V., E.I, L.K, and B.S provided critical feedback and helped shape the research, analysis and manuscript. G.Z. supervised the project.

## Appendix A GP weight map

The GP weights encode the contribution of voxels in predicting memory performance score and classifying biomarkers positivity. To compute the GP weight vector, we start by considering the posterior weight distribution (for more details see (Rasmussen, 2006) Chapter 2) given by

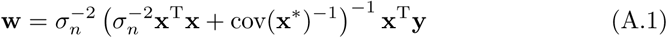

Eq. A.1 requires a huge matrix inversion, so we derivate an alternate equivalent representation, where we invert s × s matrix instead of d × d matrix (≪d, d represents number of voxels).

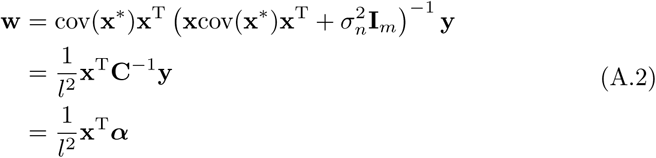

were 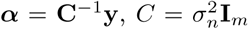. This method can be used to estimate weight contribution of each voxels for classification and regression tasks when a linear kernel is used.(Schulz et al., 2017).

## Footnotes

2 DELCODE is retrospectively registered at the German Clinical Trials Register (DRKS00007966), (04/05/2015). The study has been approved by the ethics commission and the local institutional review boards.

## References

Abdulkadir, A., Ronneberger, O., Tabrizi, S. J., and Klöppel, S. (2014). Reduction of confounding effects with voxel-wise gaussian process regression in structural mri. In 2014 International Workshop on Pattern Recognition in Neuroimaging, pages 1–4. IEEE.

Aksman, L. M., Scelsi, M. A., Marquand, A. F., Alexander, D. C., Ourselin, S., Altmann, A., and ADNI (2019). Modeling longitudinal imaging biomarkers with parametric bayesian multi-task learning. Human brain mapping, 40(13):3982–4000.

Alfaro-Almagro, F., McCarthy, P., Afyouni, S., Andersson, J. L., Bastiani, M., Miller, K. L., Nichols, T. E., and Smith, S. M. (2021). Confound modelling in uk biobank brain imaging. NeuroImage, 224:117002.

Amyot, F., Arciniegas, D. B., Brazaitis, M. P., Curley, K. C., Diaz-Arrastia, R., Gandjbakhche, A., Herscovitch, P., Hinds, S. R., Manley, G. T., Pacifico, A., et al. (2015). A review of the effectiveness of neuroimaging modalities for the detection of traumatic brain injury. Journal of neurotrauma, 32(22):1693–1721.

Ansart, M., Epelbaum, S., Gagliardi, G., Colliot, O., Dormont, D., Dubois, B., Hampel, H., Durrleman, S., Initiative, A. D. N.*, and the INSIGHT-preAD study (2020). Reduction of recruitment costs in preclinical ad trials: validation of automatic pre-screening algorithm for brain amyloidosis. Statistical methods in medical research, 29(1):151–164.

Arbabshirani, M. R., Plis, S., Sui, J., and Calhoun, V. D. (2017). Single subject prediction of brain disorders in neuroimaging: Promises and pitfalls. Neuroimage, 145:137–165.

Ashburner, J. and Friston, K. J. (2009). Computing average shaped tissue probability templates. Neuroimage, 45(2):333–341.

Ashburner, J. and Friston, K. J. (2011). Diffeomorphic registration using geodesic shooting and gauss–newton optimisation. NeuroImage, 55(3):954–967.

Ashburner, J. and Klöppel, S. (2011). Multivariate models of inter-subject anatomical variability. Neuroimage, 56(2):422–439.

Ba, M., Ng, K., Gao, X., Kong, M., Guan, L., Yu, L., and Initiative, A. D. N. (2019). The combination of apolipoprotein e4, age and alzheimer’s disease assessment scale–cognitive subscale improves the prediction of amyloid positron emission tomography status in clinically diagnosed mild cognitive impairment. European journal of neurology, 26(5):733–e53.

Bach, F. R., Lanckriet, G. R., and Jordan, M. I. (2004). Multiple kernel learning, conic duality, and the smo algorithm. In Proceedings of the twenty-first international conference on Machine learning, page 6.

Barber, N. (2005). Educational and ecological correlates of iq: A cross-national investigation. Intelligence, 33(3):273–284.

Bartrés-Faz, D. and Arenaza-Urquijo, E. M. (2011). Structural and functional imaging correlates of cognitive and brain reserve hypotheses in healthy and pathological aging. Brain topography, 24(3):340–357.

Beason-Held, L. L., Goh, J. O., An, Y., Kraut, M. A., O’Brien, R. J., Ferrucci, L., and Resnick, S. M. (2013). Changes in brain function occur years before the onset of cognitive impairment. Journal of Neuroscience, 33(46):18008–18014.

Besson, F. L., La Joie, R., Doeuvre, L., Gaubert, M., Mézenge, F., Egret, S., Landeau, B., Barré, L., Abbas, A., Ibazizene, M., et al. (2015). Cognitive and brain profiles associated with current neuroimaging biomarkers of preclinical alzheimer’s disease. Journal of Neuroscience, 35(29):10402–10411.

Bishop, C. M. (2006). Pattern recognition. Machine learning, 128(9).

Blennow, K., Zetterberg, H., and Fagan, A. M. (2012). Fluid biomarkers in alzheimer disease. Cold Spring Harbor perspectives in medicine, 2(9):a006221.

Bouwman, F., Schoonenboom, S., van Der Flier, W., Van Elk, E., Kok, A., Barkhof, F., Blankenstein, M., and Scheltens, P. (2007). Csf biomarkers and medial temporal lobe atrophy predict dementia in mild cognitive impairment. Neurobiology of aging, 28(7):1070–1074.

Bradley, R. H. and Caldwell, B. M. (1980). The relation of home environment, cognitive competence, and iq among males and females. Child Development, pages 1140–1148.

Buckley, R. F., Sikkes, S., Villemagne, V. L., Mormino, E. C., Rabin, J. S., Burnham, S., Papp, K. V., Doré, V., Masters, C. L., Properzi, M. J., et al. (2019). Using subjective cognitive decline to identify high global amyloid in community-based samples: a cross-cohort study. Alzheimer’s & Dementia: Diagnosis, Assessment & Disease Monitoring, 11(1):670–678.

Challis, E., Hurley, P., Serra, L., Bozzali, M., Oliver, S., and Cercignani, M. (2015). Gaussian process classification of alzheimer’s disease and mild cognitive impairment from resting-state fmri. NeuroImage, 112:232–243.

Chételat, G., Eustache, F., Viader, F., Sayette, V. D. L., Pélerin, A., Mézenge, F., Hannequin, D., Dupuy, B., Baron, J.-C., and Desgranges, B. (2005). Fdgpet measurement is more accurate than neuropsychological assessments to predict global cognitive deterioration in patients with mild cognitive impairment. Neurocase, 11(1):14–25.

Cummings, J., Aisen, P. S., DuBois, B., Frölich, L., Jack, C. R., Jones, R. W., Morris, J. C., Raskin, J., Dowsett, S. A., and Scheltens, P. (2016). Drug development in alzheimer’s disease: the path to 2025. Alzheimer’s research & therapy, 8(1):1–12.

Davatzikos, C., Genc, A., Xu, D., and Resnick, S. M. (2001). Voxel-based morphometry using the RAVENS maps: methods and validation using simulated longitudinal atrophy. NeuroImage, 14(6):1361–1369.

Davatzikos, C., Sotiras, A., Fan, Y., Habes, M., Erus, G., Rathore, S., Bakas, S., Chitalia, R., Gastounioti, A., and Kontos, D. (2019). Precision diagnostics based on machine learning-derived imaging signatures. Magnetic resonance imaging, 64:49–61.

Doraiswamy, P. M., Charles, H. C., and Krishnan, K. R. R. (1998). Prediction of cognitive decline in early alzheimer’s disease. The Lancet, 352(9141):1678.

Dosenbach, N. U., Nardos, B., Cohen, A. L., Fair, D. A., Power, J. D., Church, J. A., Nelson, S. M., Wig, G. S., Vogel, A. C., Lessov-Schlaggar, C. N., et al. (2010). Prediction of individual brain maturity using fmri. Science, 329(5997):1358–1361.

Dowling, N. M., Hermann, B., La Rue, A., and Sager, M. A. (2010). Latent structure and factorial invariance of a neuropsychological test battery for the study of preclinical alzheimer’s disease. Neuropsychology, 24(6):742.

Dubois, B., Hampel, H., Feldman, H. H., Scheltens, P., Aisen, P., Andrieu, S., Bakardjian, H., Benali, H., Bertram, L., Blennow, K., et al. (2016). Preclinical alzheimer’s disease: definition, natural history, and diagnostic criteria. Alzheimer’s & Dementia, 12(3):292–323.

Dubois, J., Galdi, P., Paul, L. K., and Adolphs, R. (2018). A distributed brain network predicts general intelligence from resting-state human neuroimaging data. Philosophical Transactions of the Royal Society B: Biological Sciences, 373(1756):20170284.

Duchesne, S., Caroli, A., Geroldi, C., Collins, D. L., and Frisoni, G. B. (2009). Relating one-year cognitive change in mild cognitive impairment to baseline mri features. Neuroimage, 47(4):1363–1370.

Dyrba, M., Grothe, M., Kirste, T., and Teipel, S. J. (2015). Multimodal analysis of functional and structural disconnection in a lzheimer’s disease using multiple kernel svm. Human brain mapping, 36(6):2118–2131.

Dyrba, M., Hanzig, M., Altenstein, S., Bader, S., Ballarini, T., Brosseron, F., Buerger, K., Cantré, D., Dechent, P., Dobisch, L., et al. (2021). Improving 3d convolutional neural network comprehensibility via interactive visualization of relevance maps: evaluation in alzheimer’s disease. Alzheimer’s research & therapy, 13(1):1–18.

Eustache, P., Nemmi, F., Saint-Aubert, L., Pariente, J., and Péran, P. (2016). Multimodal magnetic resonance imaging in alzheimer’s disease patients at prodromal stage. Journal of Alzheimer’s Disease, 50(4):1035–1050.

Ezzati, A., Harvey, D. J., Habeck, C., Golzar, A., Qureshi, I. A., Zammit, A. R., Hyun, J., Truelove-Hill, M., Hall, C. B., Davatzikos, C., et al. (2020). Predicting amyloid-β levels in amnestic mild cognitive impairment using machine learning techniques. Journal of Alzheimer’s Disease, 73(3):1211–1219.

Filippone, M., Marquand, A. F., Blain, C. R., Williams, S. C., Mourão-Miranda, J., and Girolami, M. (2012). Probabilhe2020deepistic prediction of neurological disorders with a statistical assessment of neuroimaging data modalities. The annals of applied statistics, 6(4):1883.

Fisher, C. K., Smith, A. M., and Walsh, J. R. (2019). Machine learning for comprehensive forecasting of alzheimer’s disease progression. Scientific reports, 9(1):1–14.

Folstein, M. F., Folstein, S. E., and McHugh, P. R. (1975). “mini-mental state”: a practical method for grading the cognitive state of patients for the clinician. Journal of psychiatric research, 12(3):189–198.

Forsberg, A., Engler, H., Almkvist, O., Blomquist, G., Hagman, G., Wall, A., Ringheim, A., Långström, B., and Nordberg, A. (2008). Pet imaging of amyloid deposition in patients with mild cognitive impairment. Neurobiology of aging, 29(10):1456–1465.

Franke, K., Ziegler, G., Klöppel, S., Gaser, C., and the Alzheimer Disease Neuroimaging Initiative (2010). Estimating the age of healthy subjects from T1-weighted MRI scans using kernel methods: exploring the influence of various parameters. NeuroImage, 50(3):883–892.

Frisoni, G. B., Fox, N. C., Jack, C. R., Scheltens, P., and Thompson, P. M. (2010). The clinical use of structural mri in alzheimer disease. Nature Reviews Neurology, 6(2):67–77.

Gaugler, J., James, B., Johnson, T., Marin, A., and Weuve, J. (2019). 2019 alzheimer’s disease facts and figures. ALZHEIMERS & DEMENTIA, 15(3):321–387.

Gönen, M. and Alpaydın, E. (2011). Multiple kernel learning algorithms. The Journal of Machine Learning Research, 12:2211–2268.

Grober, E., Ocepek-Welikson, K., and Teresi, J. A. (2009). The free and cued selective reminding test: evidence of psychometric adequacy. Psychology Science Quarterly, 51(3):266–282.

Grosenick, L., Klingenberg, B., Katovich, K., Knutson, B., and Taylor, J. E. (2013). Interpretable whole-brain prediction analysis with graphnet. NeuroImage, 72:304–321.

Gupta, Y., Lama, R. K., Kwon, G.-R., Weiner, M. W., Aisen, P., Weiner, M., Petersen, R., Jack Jr, C. R., Jagust, W., Trojanowki, J. Q., et al. (2019). Prediction and classification of alzheimer’s disease based on combined features from apolipoprotein-e genotype, cerebrospinal fluid, mr, and fdg-pet imaging biomarkers. Frontiers in computational neuroscience, 13:72.

He, T., Kong, R., Holmes, A. J., Nguyen, M., Sabuncu, M. R., Eickhoff, S. B., Bzdok, D., Feng, J., and Yeo, B. T. (2020). Deep neural networks and kernel regression achieve comparable accuracies for functional connectivity prediction of behavior and demographics. NeuroImage, 206:116276.

Hu, Y., Hosseini, A., Kauwe, J. S., Gross, J., Cairns, N. J., Goate, A. M., Fagan, A. M., Townsend, R. R., and Holtzman, D. M. (2007). Identification and validation of novel csf biomarkers for early stages of alzheimer’s disease. PROTEOMICS–Clinical Applications, 1(11):1373–1384.

Humpel, C. (2011). Identifying and validating biomarkers for alzheimer’s disease. Trends in biotechnology, 29(1):26–32.

Insel, P. S., Palmqvist, S., Mackin, R. S., Nosheny, R. L., Hansson, O., Weiner, M. W., Mattsson, N., and Initiative, A. D. N. (2016). Assessing risk for preclinical β-amyloid pathology with apoe, cognitive, and demographic information. Alzheimer’s & Dementia: Diagnosis, Assessment & Disease Monitoring, 4(1):76–84.

Jack, C. R., Bennett, D. A., Blennow, K., Carrillo, M. C., Feldman, H. H., Frisoni, G. B., Hampel, H., Jagust, W. J., Johnson, K. A., Knopman, D. S., et al. (2016). A/t/n: an unbiased descriptive classification scheme for alzheimer disease biomarkers. Neurology, 87(5):539–547.

Jack Jr, C. R., Knopman, D. S., Jagust, W. J., Petersen, R. C., Weiner, M. W., Aisen, P. S., Shaw, L. M., Vemuri, P., Wiste, H. J., Weigand, S. D., et al. (2013a). Tracking pathophysiological processes in alzheimer’s disease: an updated hypothetical model of dynamic biomarkers. The Lancet Neurology, 12(2):207–216.

Jack Jr, C. R., Knopman, D. S., Jagust, W. J., Petersen, R. C., Weiner, M. W., Aisen, P. S., Shaw, L. M., Vemuri, P., Wiste, H. J., Weigand, S. D., et al. (2013b). Update on hypothetical model of alzheimer’s disease biomarkers. Lancet neurology, 12(2):207.

Jansen, W. J., Ossenkoppele, R., Tijms, B. M., Fagan, A. M., Hansson, O., Klunk, W. E., Van Der Flier, W. M., Villemagne, V. L., Frisoni, G. B., Fleisher, A. S., et al. (2018). Association of cerebral amyloid-β aggregation with cognitive functioning in persons without dementia. JAMA psychiatry, 75(1):84–95.

Jessen, F., Amariglio, R., Boxtel, M., Breteler, M., Ceccaldi, M., Chételat, G., Dubois, B., Dufouil, C., Ellis, K., Flier, W., Glodzik, L., Harten, A. V., Leon, M., McHugh, P., Mielke, M., Molinuevo, J., Mosconi, L., Osorio, R., Perrotin, A., Petersen, R., Rabin, L., Rami, L., Reisberg, B., Rentz, D., Sachdev, P., Sayette, V., Saykin, A., Scheltens, P., Shulman, M. B., Slavin, M., Sperling, R., Stewart, R., Uspenskaya, O., Vellas, B., Visser, P., and Wagner, M. (2014). A conceptual framework for research on subjective cognitive decline in preclinical alzheimer’s disease. Alzheimer’s & Dementia, 10:844–852.

Jessen, F., Spottke, A., Boecker, H., Brosseron, F., Buerger, K., Catak, C., Fliessbach, K., Franke, C., Fuentes, M., Heneka, M. T., et al. (2018). Design and first baseline data of the dzne multicenter observational study on predementia alzheimer’s disease (delcode). Alzheimer’s research & therapy, 10(1):1–10.

Jo, T., Nho, K., and Saykin, A. J. (2019). Deep learning in alzheimer’s disease: diagnostic classification and prognostic prediction using neuroimaging data. Frontiers in aging neuroscience, 11:220.

Johnson, K. A., Fox, N. C., Sperling, R. A., and Klunk, W. E. (2012). Brain imaging in alzheimer disease. Cold Spring Harbor perspectives in medicine, 2(4):a006213.

Jollans, L., Boyle, R., Artiges, E., Banaschewski, T., Desrivières, S., Grigis, A., Martinot, J.-L., Paus, T., Smolka, M. N., Walter, H., et al. (2019). Quantifying performance of machine learning methods for neuroimaging data. NeuroImage, 199:351–365.

Kandiah, N., Zhang, A., Cenina, A. R., Au, W. L., Nadkarni, N., and Tan, L. C. (2014). Montreal cognitive assessment for the screening and prediction of cognitive decline in early parkinson’s disease. Parkinsonism & related disorders, 20(11):1145–1148.

Kim, H.-C. and Ghahramani, Z. (2006). Bayesian gaussian process classification with the em-ep algorithm. IEEE Transactions on Pattern Analysis and Machine Intelligence, 28(12):1948–1959.

Knešaurek, K. (2015). Improving 18f-fluoro-d-glucose-positron emission tomography/computed tomography imaging in alzheimer’s disease studies. World journal of nuclear medicine, 14(3):171.

Ko, H., Ihm, J.-J., Kim, H.-G., Initiative, A. D. N., et al. (2019). Cognitive profiling related to cerebral amyloid beta burden using machine learning approaches. Frontiers in aging neuroscience, 11:95.

Kohavi, R. et al. (1995). A study of cross-validation and bootstrap for accuracy estimation and model selection. In Ijcai, volume 14, pages 1137–1145. Montreal, Canada.

Korolev, I. O., Symonds, L. L., Bozoki, A. C., and Initiative, A. D. N. (2016). Predicting progression from mild cognitive impairment to alzheimer’s dementia using clinical, mri, and plasma biomarkers via probabilistic pattern classification. PloS one, 11(2):e0138866.

Lee, J. H., Byun, M. S., Yi, D., Sohn, B. K., Jeon, S. Y., Lee, Y., Lee, J.-Y., Kim, Y. K., Lee, Y.-S., and Lee, D. Y. (2018). Prediction of cerebral amyloid with common information obtained from memory clinic practice. Frontiers in aging neuroscience, 10:309.

Lezak, M. D., Howieson, D. B., Loring, D. W., Fischer, J. S., et al. (2004). Neuropsychological assessment. Oxford University Press, USA.

Lindsay, J., Laurin, D., Verreault, R., Hébert, R., Helliwell, B., Hill, G. B., and McDowell, I. (2002). Risk factors for alzheimer’s disease: a prospective analysis from the canadian study of health and aging. American journal of epidemiology, 156(5):445–453.

Liu, M., Zhang, J., Adeli, E., and Shen, D. (2018). Joint classification and regression via deep multi-task multi-channel learning for alzheimer’s disease diagnosis. IEEE Transactions on Biomedical Engineering, 66(5):1195–1206.

M Di Battista, A., M Heinsinger, N., and William Rebeck, G. (2016). Alzheimer’s disease genetic risk factor apoe-ε4 also affects normal brain function. Current Alzheimer Research, 13(11):1200–1207.

Marquand, A., Howard, M., Brammer, M., Chu, C., Coen, S., and Mourão-Miranda, J. (2010). Quantitative prediction of subjective pain intensity from whole-brain fmri data using gaussian processes. Neuroimage, 49(3):2178–2189.

Marquand, A. F., Brammer, M., Williams, S. C., and Doyle, O. M. (2014). Bayesian multi-task learning for decoding multi-subject neuroimaging data. NeuroImage, 92:298–311.

Maserejian, N., Bian, S., Wang, W., Jaeger, J., Syrjanen, J. A., Aakre, J., Jack Jr, C. R., Mielke, M. M., Gao, F., Initiative, A. D. N., et al. (2019). Practical algorithms for amyloid β probability in subjective or mild cognitive impairment. Alzheimer’s & Dementia: Diagnosis, Assessment & Disease Monitoring, 11:710–720.

Mateos-Pérez, J. M., Dadar, M., Lacalle-Aurioles, M., Iturria-Medina, Y., Zeighami, Y., and Evans, A. C. (2018). Structural neuroimaging as clinical predictor: A review of machine learning applications. NeuroImage: Clinical, 20:506–522.

McKhann, G. M., Knopman, D. S., Chertkow, H., Hyman, B. T., Jack Jr, C. R., Kawas, C. H., Klunk, W. E., Koroshetz, W. J., Manly, J. J., Mayeux, R., et al. (2011). The diagnosis of dementia due to alzheimer’s disease: recommendations from the national institute on aging-alzheimer’s association workgroups on diagnostic guidelines for alzheimer’s disease. Alzheimer’s & dementia, 7(3):263–269.

Mohs, R. C., Knopman, D., Petersen, R. C., Ferris, S. H., Ernesto, C., Grundman, M., Sano, M., Bieliauskas, L., Geldmacher, D., Clark, C., et al. (1997). Development of cognitive instruments for use in clinical trials of antidementia drugs: additions to the alzheimer’s disease assessment scale that broaden its scope. Alzheimer disease and associated disorders.

Molinuevo, J. L., Rabin, L. A., Amariglio, R., Buckley, R., Dubois, B., Ellis, K. A., Ewers, M., Hampel, H., Klöppel, S., Rami, L., Reisberg, B., Saykin, A. J., Sikkes, S., Smart, C. M., Snitz, B. E., Sperling, R., van der Flier, W. M., Wagner, M., and Jessen, F. (2017). Implementation of subjective cognitive decline criteria in research studies. Alzheimer’s & Dementia, 13(3):296–311.

Monté-Rubio, G. C., Falcón, C., Pomarol-Clotet, E., and Ashburner, J. (2018). A comparison of various mri feature types for characterizing whole brain anatomical differences using linear pattern recognition methods. NeuroImage, 178:753–768.

Morris, J. C. (2005). Dementia update 2005. Alzheimer Disease & Associated Disorders, 19(2):100–117.

Mourao-Miranda, J., Bokde, A. L., Born, C., Hampel, H., and Stetter, M. (2005). Classifying brain states and determining the discriminating activation patterns: support vector machine on functional mri data. NeuroImage, 28(4):980–995.

Murphy, M. P. and LeVine III, H. (2010). Alzheimer’s disease and the amyloid-β peptide. Journal of Alzheimer’s disease, 19(1):311–323.

Ossenkoppele, R., Schonhaut, D. R., Schöll, M., Lockhart, S. N., Ayakta, N., Baker, S. L., O’Neil, J. P., Janabi, M., Lazaris, A., Cantwell, A., et al. (2016). Tau pet patterns mirror clinical and neuroanatomical variability in alzheimer’s disease. Brain, 139(5):1551–1567.

Papp, K. V., Rentz, D. M., Orlovsky, I., Sperling, R. A., and Mormino, E. C. (2017). Optimizing the preclinical alzheimer’s cognitive composite with semantic processing: the pacc5. Alzheimer’s & Dementia: Translational Research & Clinical Interventions, 3(4):668–677.

Park, L. Q., Gross, A. L., McLaren, D. G., Pa, J., Johnson, J. K., Mitchell, M., and Manly, J. J. (2012). Confirmatory factor analysis of the adni neuropsychological battery. Brain Imaging and Behavior, 6(4):528–539.

Pereira, F., Mitchell, T., and Botvinick, M. (2009). Machine learning classifiers and fmri: a tutorial overview. Neuroimage, 45(1):S199–S209.

Petermann, F. and Lepach, A. C. (2012). Wechsler memory scale. Ed. In deutscher Übersetzung und Adaptation der WMS-IV von Davis Wechsler. Frankfurt a. M.: Pearson Assessment & Information GmbH.

Pettersson-Yeo, W., Benetti, S., Marquand, A. F., Joules, R., Catani, M., Williams, S. C., Allen, P., McGuire, P., and Mechelli, A. (2014). An empirical comparison of different approaches for combining multimodal neuroimaging data with support vector machine. Frontiers in neuroscience, 8:189.

Polcher, A., Frommann, I., Koppara, A., Wolfsgruber, S., Jessen, F., and Wagner, M. (2017). Face-name associative recognition deficits in subjective cognitive decline and mild cognitive impairment. Journal of Alzheimer’s Disease, 56(3):1185–1196.

Prestia, A., Caroli, A., Wade, S. K., Van Der Flier, W. M., Ossenkoppele, R., Van Berckel, B., Barkhof, F., Teunissen, C. E., Wall, A., Carter, S. F., et al. (2015). Prediction of ad dementia by biomarkers following the nia-aa and iwg diagnostic criteria in mci patients from three european memory clinics. Alzheimer’s & Dementia, 11(10):1191–1201.

Rajapakse, J. C., Giedd, J. N., and Rapoport, J. L. (1997). Statistical approach to segmentation of single-channel cerebral MR images. IEEE transactions on medical imaging, 16(2):176–186.

Rakotomamonjy, A., Bach, F., Canu, S., and Grandvalet, Y. (2008). Simplemkl. Journal of Machine Learning Research, 9:2491–2521.

Rao, A., Monteiro, J. M., Ashburner, J., Portugal, L., Fernandes, O., De Oliveira, L., Pereira, M., and Mourao-Miranda, J. (2015). A comparison of strategies for incorporating nuisance variables into predictive neuroimaging models. In 2015 International Workshop on Pattern Recognition in NeuroImaging, pages 61–64. IEEE.

Rao, A., Monteiro, J. M., Mourao-Miranda, J., Initiative, A. D., et al. (2017). Predictive modelling using neuroimaging data in the presence of confounds. NeuroImage, 150:23–49.

Rasmussen, C. (2006). Advances in Gaussian processes. Advances in Neural Information Processing ….

Rasmussen, C. E. and Williams, C. K. I. (2006). Gaussian Processes for Machine Learning. MIT Press, Cambridge.

Rathore, S., Habes, M., Iftikhar, M. A., Shacklett, A., and Davatzikos, C. (2017). A review on neuroimaging-based classification studies and associated feature extraction methods for alzheimer’s disease and its prodromal stages. NeuroImage, 155:530–548.

Reitan, R. M. (1958). Validity of the trail making test as an indicator of organic brain damage. Perceptual and motor skills, 8(3):271–276.

Riedel, B. C., Thompson, P. M., and Brinton, R. D. (2016). Age, apoe and sex: triad of risk of alzheimer’s disease. The Journal of steroid biochemistry and molecular biology, 160:134–147.

Rohde, T. E. and Thompson, L. A. (2007). Predicting academic achievement with cognitive ability. Intelligence, 35(1):83–92.

Rouleau, I., Salmon, D. P., Butters, N., Kennedy, C., and McGuire, K. (1992). Quantitative and qualitative analyses of clock drawings in alzheimer’s and huntington’s disease. Brain and cognition, 18(1):70–87.

Salvatore, C., Battista, P., and Castiglioni, I. (2016). Frontiers for the early diagnosis of ad by means of mri brain imaging and support vector machines. Current Alzheimer Research, 13(5):509–533.

Sanderman, R., Coyne, J. C., and Ranchor, A. V. (2006). Age: Nuisance variable to be eliminated with statistical control or important concern? Patient Education and Counseling, 61(3):315–316.

Scheinost, D., Noble, S., Horien, C., Greene, A. S., Lake, E. M., Salehi, M., Gao, S., Shen, X., O’Connor, D., Barron, D. S., et al. (2019). Ten simple rules for predictive modeling of individual differences in neuroimaging. NeuroImage, 193:35–45.

Schulz, E., Speekenbrink, M., and Krause, A. (2017). A tutorial on gaussian process regression: Modelling, exploring, and exploiting functions. bioRxiv.

Shawe-Taylor, J. and Cristianini, N. (2004). Kernel methods for pattern analysis. page 462.

Shawe-Taylor, J., Cristianini, N., et al. (2004). Kernel methods for pattern analysis. Cambridge university press.

Smith, A. (1982). Symbol digit modalities test (sdmt) manual (revised) western psychological services. Los Angeles.

Stern, Y., Arenaza-Urquijo, E. M., Bartrés-Faz, D., Belleville, S., Cantilon, M., Chetelat, G., Ewers, M., Franzmeier, N., Kempermann, G., Kremen, W. S., et al. (2020). Whitepaper: Defining and investigating cognitive reserve, brain reserve, and brain maintenance. Alzheimer’s & Dementia, 16(9):1305–1311.

Stonnington, C. M., Chu, C., Klöppel, S., Jack Jr, C. R., Ashburner, J., Frackowiak, R. S., Initiative, A. D. N., et al. (2010). Predicting clinical scores from magnetic resonance scans in alzheimer’s disease. Neuroimage, 51(4):1405–1413.

Sui, J., Adali, T., Yu, Q., Chen, J., and Calhoun, V. D. (2012). A review of multivariate methods for multimodal fusion of brain imaging data. Journal of neuroscience methods, 204(1):68–81.

Thalmann, B., Monsch, A. U., Schneitter, M., Bernasconi, F., Aebi, C., Camachova-Davet, Z., and Staehelin, H. B. (2000). The cerad neuropsychological assessment battery (cerad-nab)—a minimal data set as a common tool for german-speaking europe. Neurobiology of Aging, (21):30.

Tohka, J., Zijdenbos, A., and Evans, A. (2004). Fast and robust parameter estimation for statistical partial volume models in brain MRI. NeuroImage, 23(1):84–97.

Tosun, D., Chen, Y.-F., Yu, P., Sundell, K. L., Suhy, J., Siemers, E., Schwarz, A. J., Weiner, M. W., Initiative, A. D. N., et al. (2016). Amyloid status imputed from a multimodal classifier including structural mri distinguishes progressors from nonprogressors in a mild alzheimer’s disease clinical trial cohort. Alzheimer’s & Dementia, 12(9):977–986.

Tosun, D., Joshi, S., Weiner, M. W., and Initiative, A. D. N. (2013). Neuroimaging predictors of brain amyloidosis in mild cognitive impairment. Annals of neurology, 74(2):188–198.

Van Dam, N. T., Sano, M., Mitsis, E. M., Grossman, H. T., Gu, X., Park, Y., Hof, P. R., and Fan, J. (2013). Functional neural correlates of attentional deficits in amnestic mild cognitive impairment. PLoS One, 8(1):e54035.

Varoquaux, G. (2018). Cross-validation failure: small sample sizes lead to large error bars. Neuroimage, 180:68–77.

Wang, Y., Fan, Y., Bhatt, P., and Davatzikos, C. (2010). High-dimensional pattern regression using machine learning: from medical images to continuous clinical variables. Neuroimage, 50(4):1519–1535.

Wilson, A. and Adams, R. (2013). Gaussian process kernels for pattern discovery and extrapolation. In International conference on machine learning, pages 1067–1075. PMLR.

Wolfsgruber, S., Kleineidam, L., Guski, J., Polcher, A., Frommann, I., Roeske, S., Spruth, E. J., Franke, C., Priller, J., Kilimann, I., Teipel, S., Buerger, K., Janowitz, D., Laske, C., Buchmann, M., Peters, O., Menne, F., Fuentes Casan, M., Wiltfang, J., Bartels, C., Düzel, E., Metzger, C., Glanz, W., Thelen, M., Spottke, A., Ramirez, A., Kofler, B., Fließbach, K., Schneider, A., Heneka, M. T., Brosseron, F., Meiberth, D., Jessen, F., Wagner, M., and on behalf of the DELCODE Study Group (2020). Minor neuropsychological deficits in patients with subjective cognitive decline. Neurology, 95(9):e1134–e1143.

Wolfsgruber, S., Wagner, M., Schmidtke, K., Frölich, L., Kurz, A., Schulz, S., Hampel, H., Heuser, I., Peters, O., Reischies, F. M., et al. (2014). Memory concerns, memory performance and risk of dementia in patients with mild cognitive impairment. PloS one, 9(7):e100812.

Woodard, J. L., Seidenberg, M., Nielson, K. A., Smith, J. C., Antuono, P., Durgerian, S., Guidotti, L., Zhang, Q., Butts, A., Hantke, N., et al. (2010). Prediction of cognitive decline in healthy older adults using fmri. Journal of Alzheimer’s Disease, 21(3):871–885.

Zhang, D., Shen, D., Initiative, A. D. N., et al. (2012). Multi-modal multi-task learning for joint prediction of multiple regression and classification variables in alzheimer’s disease. NeuroImage, 59(2):895–907.

Zhang, D., Wang, Y., Zhou, L., Yuan, H., Shen, D., Initiative, A. D. N., et al. (2011). Multimodal classification of alzheimer’s disease and mild cognitive impairment. Neuroimage, 55(3):856–867.

Ziegler, G., Ridgway, G. R., Dahnke, R., Gaser, C., Initiative, A. D. N., et al. (2014). Individualized gaussian process-based prediction and detection of local and global gray matter abnormalities in elderly subjects. NeuroImage, 97:333–348.

Zu, C., Jie, B., Liu, M., Chen, S., Shen, D., and Zhang, D. (2016). Label-aligned multi-task feature learning for multimodal classification of alzheimer’s disease and mild cognitive impairment. Brain imaging and behavior, 10(4):1148–1159.

